# Anxiety-related frontocortical activity is associated with dampened stressor reactivity in the real world

**DOI:** 10.1101/2021.03.17.435791

**Authors:** Juyoen Hur, Manuel Kuhn, Shannon E. Grogans, Allegra S. Anderson, Samiha Islam, Hyung Cho Kim, Rachael M. Tillman, Andrew S. Fox, Jason F. Smith, Kathryn A. DeYoung, Alexander J. Shackman

## Abstract

Negative affect is a fundamental dimension of human emotion. When extreme, it contributes to a variety of adverse outcomes—from physical and mental illness to divorce and premature death. Mechanistic work in animals and neuroimaging research in humans and monkeys has begun to reveal the broad contours of the neural circuits governing negative affect, but the relevance of these discoveries to everyday distress remains incompletely understood. Here we used a combination of approaches— including neuroimaging assays of threat anticipation and emotional face perception and >10,000 momentary assessments of emotional experience—to demonstrate that individuals showing greater activation in a cingulo-opercular circuit during an anxiety-eliciting laboratory paradigm experience lower levels of stressor-dependent distress in their daily lives (*n*=202-208). Extended amygdala activation was not significantly related to momentary negative affect. These observations provide a framework for understanding the neurobiology of negative affect in the laboratory and in the real world.

**STATEMENT OF RELEVANCE:** Anxiety, sadness, and other negative emotions are hallmarks of the human condition. When extreme, they contribute to a variety of adverse outcomes—from physical and mental illness to divorce and premature death—pointing to the need to develop a better understanding of the underlying brain circuitry. Recent work has begun to reveal the neural systems governing negative affect, but the relevance of these tantalizing laboratory discoveries to the real world has remained unclear. Here we used a combination of brain imaging and smartphone-based survey techniques to show that individuals marked by greater activation in a cingulo-opercular circuit during an anxiety-promoting laboratory paradigm tend to experience diminished distress in response to everyday stressors. These observations provide new insights into the brain systems most relevant to moment-by-moment fluctuations in negative mood, underscoring the importance of more recently evolved cortical association areas.

## INTRODUCTION

Negative affect is a fundamental dimension of mammalian emotion. It encompasses transient states—like anxiety, fear, sadness, and worry—and more persistent, trait-like tendencies to experience and express negative emotions (Shackman et al., 2016). When extreme or pervasive, negative affect contributes to a panoply of adverse outcomes—from physical and mental illness to divorce and premature death— underscoring the need to develop a better understanding of the underlying neurobiology (Hur, Stockbridge, Fox, & Shackman, 2019).

Mechanistic work in animals and neuroimaging research in humans and monkeys has begun to reveal the broad contours of the neural systems governing negative affect (Chang, Gianaros, Manuck, Krishnan, & Wager, 2015; Fox & Shackman, 2019; Horikawa, Cowen, Keltner, & Kamitani, 2020; Kenwood & Kalin, 2021; Taschereau-Dumouchel, Kawato, & Lau, 2019). This work underscores the importance of *subcortical* regions, like the amygdala, bed nucleus of the stria terminalis (BST), and periaqueductal gray (PAG). But it also highlights *frontocortical* regions that are particularly well developed in humans, including the midcingulate cortex (MCC), anterior insula (AI), frontal operculum (FrO), and dorsolateral prefrontal cortex (dlPFC) (Hur et al., 2020b; Shackman & Fox, 2021; Shackman et al., 2011). At present, the relevance of these tantalizing laboratory discoveries to subjective emotional experience in the real world remains incompletely understood. Given the limitations of ambulatory measures of brain activity, overcoming this barrier requires integrating measures of emotion-relevant brain function acquired in the laboratory with assessments of negative affect collected in the field.

Here we used fMRI to quantify individual differences in neural reactivity to a well-established anxiety-provocation (‘threat-anticipation’) paradigm in 220 young adults (**Figure 1**). A multiband MRI sequence and best-practice data processing techniques enhanced our ability to resolve small subcortical regions (e.g., amygdala and BST). Next, we used smartphone experience sampling—often termed ‘ecological momentary assessment’ (EMA)—to intensively sample fluctuations in self-reported negative affect and stressor exposure across different real-world contexts. Because EMA data are captured in real time, they circumvent the biases that can distort retrospective reports and provide insights into how emotional experience dynamically responds to everyday stressors (Lay, Gerstorf, Scott, Pauly, & Hoppmann, 2017; Shiffman, Stone, & Hufford, 2008). To ensure a broad spectrum of emotional reactivity, subjects were selectively recruited from a pool of 6,594 young adults screened for trait-like individual differences in negative emotionality, consistent with recent recommendations (Charpentier et al., *in press*). We focused on ‘emerging adulthood’ because it is a time of profound, often stressful transitions (Shackman et al., 2018). In fact, more than half of undergraduate students report moderate-to-severe levels of anxiety and depression, with many experiencing frank emotional disorders during this turbulent developmental chapter (NASEM, 2021; SAMHSA, 2019; Vos et al., 2020).

**Figure 1.**
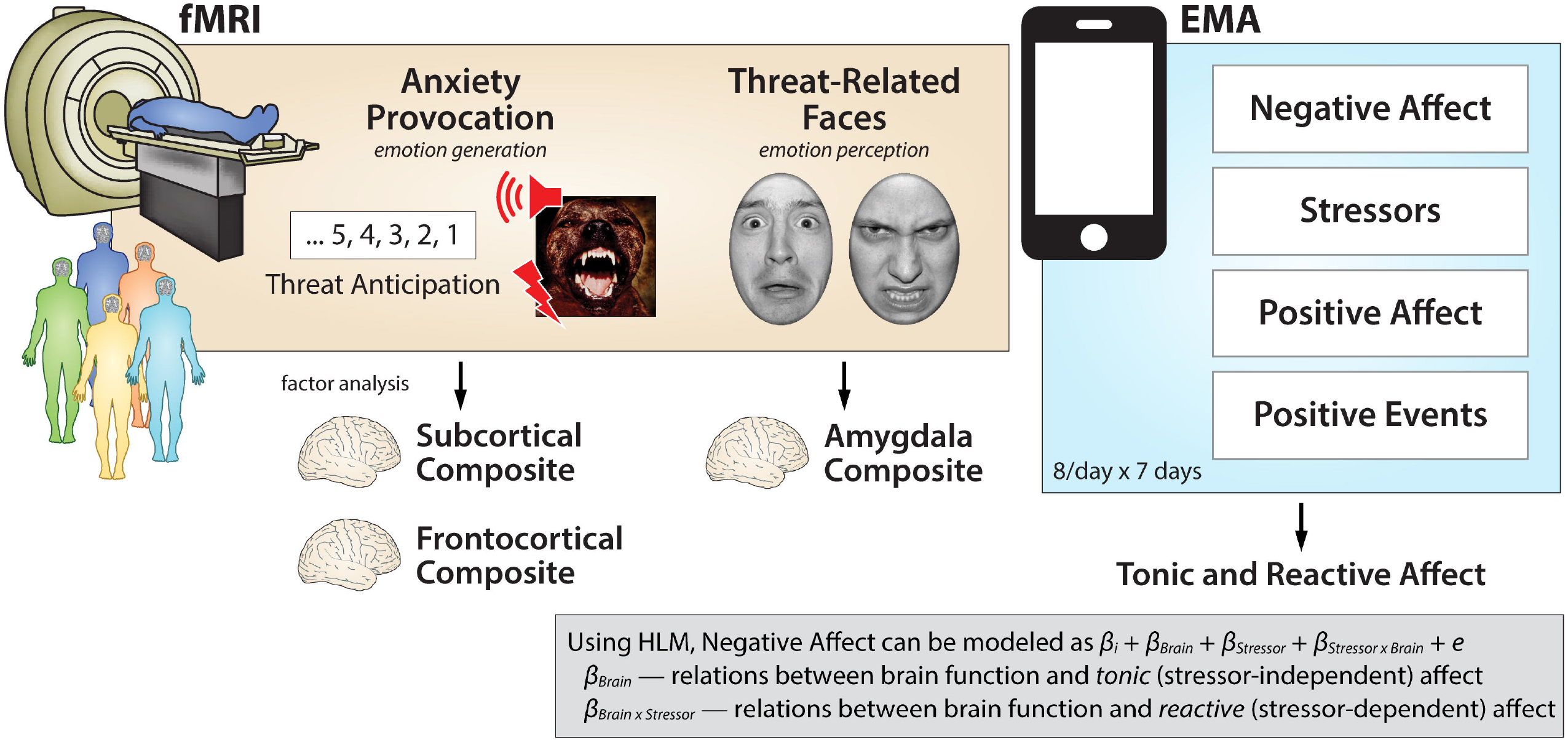
Overview of fMRI-EMA fusion. fMRI. All subjects were assessed using fMRI and a well-established anxiety-provocation paradigm. On Threat trials, subjects saw a stream of integers that culminated with the delivery of a noxious electric shock, unpleasant photograph, and thematically related audio clip (e.g., growl, scream). Control trials were similar, but terminated with the delivery of benign reinforcers. A subset of subjects also completed a threat-related faces paradigm, during which they viewed photographs of fearful or angry faces. Photographs of neutral outdoor scenes served as a control. To integrate the fMRI and EMA data-streams, regions of interest were prescribed based on significant group-level activation in whole-brain analyses (FDR *q* < 0.05, whole-brain corrected) within anatomical regions highlighted by prior work using similar tasks. For the anxiety-provocation task, analyses focused on *subcortical* (amygdala, BST, and PAG) and *frontocortical* (MCC, AI, FrO, and dlPFC) activation during threat anticipation. To reduce the number of comparisons and maximize statistical power, a factor analysis was used to guide the construction of composite measures of brain activity. For the faces paradigm, analyses focused on a composite measure of bilateral amygdala activation. **EMA.** Smartphone EMA was used to sample hour-by-hour fluctuations in negative and positive affect. Subjects also reported exposure to stressors and positive events in the past hour, enabling examination of *tonic* (contextindependent) and *reactive* (context-dependent) affect. Subjects completed up to 8 EMA surveys per day for 7 days, yielding a total of 10,239 usable assessments. **fMRI-EMA fusion.** A series of hierarchical linear models (HLMs) enabled us to selectively probe relations between the composite neuroimaging metrics derived for each task and tonic and reactive negative affect. A similar approach was used for follow-up tests of specificity and incremental validity. Abbreviation—*e*, error (unmodeled variance); EMA, ecological momentary assessment; fMRI, functional magnetic resonance imaging; HLM, hierarchical linear modeling; *i*, intercept.

A series of hierarchical linear models (HLMs)—sometimes termed ‘multilevel’ or ‘linear mixed’ models— was used to fuse the fMRI and EMA data-streams (**Figure 1**). HLM naturally handles the nested dependency and variable number of assessments completed by each subject, is the standard analytic framework for EMA and other kinds of experience-sampling data (Shiffman et al., 2008), and enabled us to test relations between anxiety-related brain function and *tonic* (stressor-independent) and *reactive* (stressor-dependent) variation in real-world negative affect (Gross, Sutton, & Ketelaar, 1998; Shackman et al., 2016). To clarify specificity, parallel analyses were performed for positive affect and positive events. This analytic framework also allowed us to quantify the added explanatory value (‘incremental validity’) of neuroimaging metrics relative to conventional paper-and-pencil measures of trait negative emotionality (Shackman & Fox, 2018).

Aside from addressing the real-world significance of anxiety-related brain circuitry, this approach afforded an opportunity to clarify the contributions of frontocortical regions to negative affect. Although the MCC, AI/FrO, and dlPFC are consistently recruited by a variety of distress-eliciting experimental challenges, their precise role has remained enigmatic (Hur et al., 2020b). In part, this reflects the fact that a broadly similar network is recruited by emotion-regulation paradigms (Langner, Leiberg, Hoffstaedter, & Eickhoff, 2018; Morawetz et al., 2020), raising the possibility that frontocortical activation actually reflects spontaneous efforts to dampen, rather than promote, distress. Fusing the fMRI and EMA data streams enabled us to test whether frontocortical activity is associated with increased or decreased negative affect in the midst of daily life.

To provide a more direct link with on-going research, parallel analyses were conducted using fMRI data from a subset of subjects who also completed an emotional-faces paradigm. Variants of the emotional-faces paradigm are widely used as probes of amygdala function—often in the guise of quantifying variation in ‘Negative Valence Systems’—and have been incorporated into many biobank studies, including ABCD (Casey et al., 2018), the Human Connectome Project (HCP) and follow-ons (e.g., Barch et al., 2013; Seok et al., 2020), IMAGEN (Albaugh et al., 2019), the Philadelphia Neurodevelopmental Cohort (Satterthwaite et al., 2016), and UK Biobank (Miller et al., 2016). Although photographs of models posing ‘threat-related’ (i.e., fearful and angry) facial expressions strongly activate the amygdala (Miller et al., 2016), they do not elicit distress in typical adults and, as such, are better conceptualized as an assay of emotion perception, rather than the experience or expression of emotion (Hur et al., 2019). Here, we tested whether differences in amygdala reactivity to threat-related faces are associated with EMA measures of negative affect (**Figure 1**).

Discovering the neural systems most relevant to the moment-by-moment experience of negative affect in daily life is important. Emotional illnesses are defined, diagnosed, and treated on the basis of real-world feelings, and for some theorists they are *the* defining feature of emotion (Fox, Lapate, Shackman, & Davidson, 2018; LeDoux & Pine, 2016; Mobbs et al., 2019). This approach has the potential to provide insights that cannot be achieved using either animal models or isolated measures of human brain function and represents a step to establishing the everyday relevance of the brain circuits highlighted in neuroimaging studies of emotion perception and generation.

## METHOD

### Overview

As part of an on-going prospective-longitudinal study focused on the emergence of internalizing disorders, we used measures of trait negative emotionality—often termed neuroticism or dispositional negativity—to screen 6,594 young adults (57.1% female, 42.9% male; 59.0% White, 19.0% Asian, 9.9% African American, 6.3% Hispanic, 5.8% Multiracial/Other; *M* = 19.2 years, *SD* = 1.1 years) (Hur et al., 2020a; Shackman et al., 2018). Screening data were stratified into quartiles (top quartile, middle quartiles, bottom quartile) separately for men and women. Individuals who met preliminary inclusion criteria were independently and randomly recruited from each of the resulting six strata. Given the focus of the larger study, approximately half the subjects were recruited from the top quartile, with the remainder split between the middle and bottom quartiles (i.e., 50% high, 25% medium, and 25% low), enabling us to sample a wide range of risk for the development of internalizing disorders. Simulation work suggests that this enrichment approach does not bias statistical tests to a degree that would compromise their validity (Hauner, Zinbarg, & Revelle, 2014). All subjects were first-year university students in good physical health with normal or corrected-to-normal color vision and access to a personal smartphone. All reported the absence of lifetime neurological or pervasive developmental disorders, MRI contraindications, or prior experience with aversive electrical stimulation. All subjects were free from lifetime psychotic and bipolar disorders; a current mood, anxiety, or trauma disorder (past 2 months); severe substance abuse (i.e., associated with physical disability, hospitalization, or inpatient treatment); active suicidality; and on-going psychiatric treatment as determined by an experienced masters-level diagnostician using the Structured Clinical Interview for DSM-5 (First, Williams, Karg, & Spitzer, 2015). To maximize the range of emotional reactivity and psychiatric risk, subjects with a lifetime history of internalizing disorders were not excluded. At the baseline laboratory session, subjects provided informed written consent, were familiarized with the EMA protocol (see below), and re-completed the assessment of trait negative emotionality. Beginning the next day, subjects completed up to 8 EMA surveys per day for 7 days (see below). Subjects completed the neuroimaging assessment within 2-5 weeks of beginning the EMA protocol (*Median* = 13 days, *Max* = 39 days). All procedures were approved by the University of Maryland Institutional Review Board. The sample overlaps that featured in work by our group focused on social anxiety and momentary mood (Hur et al., 2020a) and the basic neurobiology of threat processing (Hur et al., 2020b).

### Subjects

A total of 234 subjects completed the MRI assessment.

#### Anxiety Provocation

Fourteen subjects were excluded from fMRI analyses of the anxiety-provocation task due to incidental neurological findings (*n* = 4), scanner problems (*n* = 2), insufficient usable fMRI data (*n* = 2; see below), or excessive global motion artifact (*n* = 6; see below). Twelve subjects did not successfully complete the EMA assessment (see below), and were excluded from fMRI-EMA analyses. This yielded a final sample of 208 subjects (50.0% female, 50.0% male; 63.0% White, 17.3% Asian, 8.2% African American, 3.8% Hispanic, 7.7% Multiracial/Other; *M* = 18.8 years, *SD* = 0.4).

#### Threat-Related Faces

Twenty-one subjects were excluded from fMRI analyses of the threat-related faces task due to incidental neurological findings (*n* = 4), scanner problems (*n* = 2), gross artifacts (*n* = 1), insufficient usable fMRI data (*n* = 1; see below), excessive global motion artifact (*n* = 7; see below), or inadequate performance accuracy (*n* = 6; see below). Eleven subjects did not successfully complete the EMA assessment (see below), and were excluded from fMRI-EMA analyses. This yielded a final sample of 202 subjects (50.0% female, 50.0% male; 63.4% White, 16.8% Asian, 8.4% African American, 4.0% Hispanic, 7.4% Multiracial/Other; *M* = 18.8 years, *SD* = 0.4).

### Power Analysis

Sample size was determined *a priori* as part of the application for the award that supported data collection (R01-MH107444). The target sample size (*N* ≈ 240) was chosen to afford acceptable power and precision given available resources (Schönbrodt & Perugini, 2013). At the time of study design, G-power 3.1.9.2 (http://www.gpower.hhu.de) indicated >99% power to detect a benchmark medium-sized effect (*r* = .30) with up to 20%planned attrition (*N* = 192 usable datasets) using α = .05 (two-tailed).

### Trait Negative Emotionality

As in prior work (Hur et al., 2020a; Shackman et al., 2018), we used measures of neuroticism (Big Five Inventory-Neuroticism, α = 0.86-0.89; John, Naumann, & Soto, 2008) and trait anxiety (International Personality Item Pool-Trait Anxiety, α = 0.89; Goldberg, 1999; Goldberg et al., 2006) to quantify stable individual differences in negative emotionality. Subjects used a 1 (*disagree strongly*) to 5 (*agree strongly*) scale to rate themselves on a total of 18 items (e.g., *depressed or blue, tense, worry, nervous, get distressed easily, fear for the worst, afraid of many things*). To minimize the influence of occasion-specific fluctuations in responding (Mõttus et al., 2020), analyses employed a measure averaged across the screening and baseline assessments (median interval = 80.0 days, *SD* = 56.6). The resulting multi-scale, multi-occasion composite captured a sizable spectrum of trait negative emotionality (*z* = −2.11 to 1.49) and showed acceptable reliability (α = 0.93, *r_Retest_* = .75).

### General Neuroimaging Procedures

Prior to scanning, subjects practiced abbreviated versions of the tasks until staff confirmed understanding. A detailed description of the peripheral apparatus is available (Hur et al., 2020b). During imaging, foam inserts were used to mitigate potential motion artifacts. Subjects were continuously monitored using an MRI-compatible eye-tracker (Eyelink 1000; SR Research, Ottawa, Ontario, Canada) and a near real-time head-motion monitor (Cox, Jesmanowicz, & Hyde, 1995). Following the last scan, subjects were removed from the scanner, debriefed, compensated, and discharged.

### Anxiety-Provocation Paradigm

#### Paradigm Structure and Procedures

The anxiety-provocation (threat-anticipation) paradigm is schematically depicted in **Supplementary Figure S1**. A detailed description of the paradigm and associated procedures and stimuli is available (Hur et al., 2020b). Subjects were informed about the task design and contingencies prior to scanning. The task was administered in 3 scans (12 trials/valence/scan). On Threat trials, subjects saw a stream of integers (*M* = 18.75 s; 8.75 - 30.00 s). To ensure robust anticipatory anxiety, this epoch always culminated with the delivery of a noxious electric shock, unpleasant photograph (e.g., mutilated body), and thematically related audio clip (e.g., scream, gunshot). Safety trials were similar but terminated with the delivery of benign reinforcers (i.e., just-perceptible electrical stimulation and neutral audiovisual stimuli). Valence was continuously signaled during the anticipation epoch by the background color of the display. White-noise visual masks (3.2 s) were presented between trials to minimize persistence of the visual reinforcers in iconic memory. Subjects were periodically prompted to rate the intensity of negative affect (‘fear/anxiety’) experienced a few seconds earlier, during the *anticipation* period of the prior trial, using a 1 (*minimal*) to 4 (*maximal*) scale.

#### Skin Conductance Data Acquisition and Processing Pipeline

To confirm the validity of the anxiety-provocation paradigm, skin conductance was continuously assessed during each scan of the anxiety-provocation task using a Biopac system (MP-150; Biopac Systems, Inc., Goleta, CA). Skin conductance (250 Hz; 0.05 Hz high-pass) was measured using MRI-compatible disposable electrodes (EL507) attached to the second and third fingers of the non-dominant hand. Skin conductance data were processed using PsPM (version 4.0.2) and in-house MATLAB code (Bach et al., 2018; Bach & Friston, 2013). Data from each scan were band-pass filtered (0.01-0.25 Hz), resampled to match the TR used for fMRI data acquisition (1.25 s), and *z*-transformed.

### Threat-Related Faces Paradigm

The threat-related faces paradigm is schematically depicted in **Supplementary Figure S2**. The paradigm took the form of a pseudo-randomized block design and was administered in 2 scans, with a short break between scans. During each scan, subjects viewed photographs of models depicting prototypical angry faces, fearful faces, happy faces, or places (7 blocks/condition/scan). Blocks consisted of 10 photographs (1.6 s) separated by fixation crosses (0.4 s). To ensure engagement, subjects judged whether the current photograph matched that presented on the prior trial (i.e., a ‘1-back’ continuous performance task). Place stimuli consisted of photographs of outdoor scenes focused on single-family residential buildings (‘houses’) or urban commercial buildings (‘skyscrapers’), and were adapted from prior work (Choi, Padmala, & Pessoa, 2012, 2015). A total of 6 additional EPI volumes were acquired at the beginning and end of each scan (see below). Subjects showing inadequate performance (i.e., accuracy <2 *SD* for both scans) were excluded from analyses.

### MRI Data Acquisition

MRI data were acquired using a Siemens Magnetom TIM Trio 3 Tesla scanner (32-channel head-coil). Sagittal T1-weighted anatomical images were acquired using a magnetization prepared rapid acquisition gradient echo (MPRAGE) sequence (TR = 2,400 ms; TE = 2.01 ms; inversion time = 1060 ms; flip angle = 8°; sagittal slice thickness = 0.8 mm; in-plane = 0.8 × 0.8 mm; matrix = 300 × 320; field-of-view = 240 × 256). A T2-weighted image was collected co-planar to the T1-weighted image (TR = 3,200 ms; TE = 564 ms; flip angle = 120°). To enhance resolution, we used a multi-band sequence to collect echo planar imaging (EPI) volumes (multiband acceleration = 6; TR = 1,250 ms; TE = 39.4 ms; flip angle = 36.4°; slice thickness = 2.2 mm, number of slices = 60; in-plane resolution = 2.1875 × 2.1875 mm; matrix = 96 × 96). Images were collected in the oblique axial plane (approximately −20° relative to the AC-PC plane) to minimize potential susceptibility artifacts (anxiety provocation: 478 volumes/scan; faces: 454 volumes/scan). The first 7 volumes were automatically discarded by the scanner. To allow field map correction, two oblique-axial spin echo (SE) images were collected in opposing phase-encoding directions (rostral-to-caudal and caudal-to-rostral) at the same location and resolution as the EPI volumes (TR = 7,220 ms, TE = 73 ms).

### MRI Data Processing Pipeline

Methods were optimized to minimize spatial normalization error and other potential sources of noise. Structural and functional MRI data were visually inspected before and after processing for quality assurance.

#### Anatomical Data

Methods were similar to those employed in recent reports by our group (e.g., Hur et al., 2020b). T1-weighted images were inhomogeneity corrected using N4 (Tustison et al., 2010)and filtered using the denoise function in ANTS (Avants et al., 2011). The brain was then extracted using a variant of BEaST (Eskildsen et al., 2012) with brain-extracted and normalized reference brains from the IXI database (https://brain-development.org/ixi-dataset). Brain-extracted T1 images were normalized to a version of the brain-extracted 1-mm T1-weighted MNI152 (version 6) template (Grabner et al., 2006) customized to remove extracerebral tissue. This was motivated by evidence that brain-extracted T1 images and templates enhance the quality of spatial normalization (Acosta-Cabronero, Williams, Pereira, Pengas, & Nestor, 2008; Fein et al., 2006; Fischmeister et al., 2013). Normalization was performed using the diffeomorphic approach implemented in SyN (version 1.9.x.2017-09.11; Avants et al., 2011; Klein et al., 2009). T2-weighted images were rigidly co-registered with the corresponding T1 prior to normalization and the brain extraction mask from the T1 was applied. Tissue priors (Lorio et al., 2016) were unwarped to the native space of each T1 using the inverse of the diffeomorphic transformation. Brain-extracted T1 and T2 images were simultaneously segmented using native-space priors generated using FAST (FSL version 5.0.9; Zhang, Brady, & Smith, 2001) for use in T1-EPI co-registration (see below).

#### Field Map Data

SE images were used to create a field map in topup (Andersson, Skare, & Ashburner, 2003; Graham, Drobnjak, & Zhang, 2017; Smith et al., 2004). Field maps were converted to radians, median filtered, and smoothed (2-mm). The average of the distortion-corrected SE images was inhomogeneity-corrected using N4 and brain-masked using 3dSkullStrip in AFNI (version 17.2.10; Cox, 1996). The resulting mask was minimally eroded to exclude extracerebral voxels.

#### Functional Data

EPI files were de-spiked (3dDespike) and slice-time corrected (to the center of the TR) using 3dTshift, inhomogeneity corrected using N4, and motion corrected to the first volume using a 12-parameter affine transformation implemented in ANTs. Recent work indicates that de-spiking is more effective than ‘scrubbing’ for attenuating motion-related artifacts (Jo et al., 2013; Power, Schlaggar, & Petersen, 2015; Siegel et al., 2014). Transformations were saved in ITK-compatible format for subsequent use. The first volume was extracted for EPI-T1 co-registration. The reference EPI volume was simultaneously co-registered with the corresponding T1-weighted image in native space and corrected for geometric distortions using boundary-based registration (Greve & Fischl, 2009). This step incorporated the previously created field map, undistorted SE, T1, white matter (WM) image, and masks. The spatial transformations necessary to transform each EPI volume from native space to the reference EPI, from the reference EPI to the T1, and from the T1 to the template were concatenated and applied to the processed (de-spiked and slice-time corrected) EPI data in a single step to minimize incidental spatial blurring. Normalized EPI data were resampled to 2-mm isotopic voxels using fifth-order b-splines and smoothed (6-mm FWHM) using 3DblurInMask.

### EMA Protocol, Measures, and Data Reduction

#### Protocol

As in other work by our group (Doorley et al., 2021; Doorley, Volgenau, Kelso, Kashdan, & Shackman, 2020; Hur et al., 2020a; Shackman et al., 2018), SurveySignal was used to automatically deliver 8 text messages/day to each subject’s smartphone (Hofmann & Patel, 2015). Messages were delivered between 8:30 AM and 11:00 PM, with 1.5-3 hours between successive messages (*M* = 120 minutes, *SD* = .43). During weekdays, messages were delivered during the ‘passing periods’ between scheduled university courses to reduce burden and maximize compliance. Messages were delivered according to a fixed schedule that varied across days (e.g., the third message was delivered at 12:52 PM on Mondays and 12:16 PM on Tuesdays). Messages contained a link to a secure on-line survey. Subjects were instructed to respond within 30 minutes (*Median* = 2 min, *SD* = 7 min) and to refrain from responding at unsafe or inconvenient moments (e.g., while driving). During the baseline laboratory session, several procedures were used to promote compliance (Palmier-Claus et al., 2011) including: (a) delivering a test message to the subject’s phone and confirming that they were able to successfully complete the on-line survey, (b) providing subjects with a 24/7 technical support number, and (c) providing monetary bonuses for increased compliance.

#### EMA Survey and Data Reduction

Current negative affect (*afraid, nervous, worried, hopeless, sad*) and positive affect (*cheerful, content, enthusiastic, joy, relaxed, calm*) at the moment of the survey prompt was rated using a 0 (*not at all*) to 4 (*extremely*) scale. Stressor exposure was assessed using a binary item (*Did you experience one or more negative events in the past hour?)*. A parallel item was used to assess recent positive events. A factor analysis (principal components extraction; oblimin rotation) of the negative and positive affect items yielded a two-factor solution, with robust loadings on the target scales (*λ*s = .64-.88) and negligible cross-loadings (*λ*s < |.19|). Both scales demonstrated adequate internal-consistency reliability at the between-subject (*α*s = .80-.88) and within-subject levels (*α*s = .92-93). Within-subject reliability was evaluated using two-level unconditional models (items nested within subjects) implemented in the *semTools* package for *R*. The reliability of the Level-1 intercept represents McDonald’s *w* adjusted for differences between subjects (Nezlek, 2007; Raudenbush & Bryk, 2002). To better understand the nature of significant brain-EMA relations, follow-up tests employed composite anxiety (*afraid, nervous, worried*) and depression (*hopeless, sad*) facet scales (*α*s = .66-.79). EMA compliance was acceptable (*M* = 87.9%, *SD* = 6.1%, Minimum = 71.4%, Total assessments = 10,239).

### Skin Conductance Modeling

Using standard MATLAB functions, SCR data were modeled using an approach similar to that used for the fMRI data. A general linear model (GLM) was used to estimate skin conductance levels during the anticipatory epoch of each condition of the anxiety-provocation paradigm for each subject (Bach, 2014; Bach, Flandin, Friston, & Dolan, 2009; Bach, Friston, & Dolan, 2013). Predictors from the first-level fMRI model (see below) were convolved with a canonical skin conductance response function (Bach, Flandin, Friston, & Dolan, 2010; Gerster, Namer, Elam, & Bach, 2018), bandpass filtered to match the data, and *z*-transformed.

### fMRI Modeling and Data Reduction

#### Data Exclusions

To assess residual global motion artifact, we computed average volume-to-volume displacement for each scan using the motion-corrected data. Scans with excess artifact (>2 *SD*) were discarded. Subjects who lacked sufficient usable fMRI data (<2 scans per task) or showed inadequate performance on the emotional-faces task (see above; accuracy <2 *SD*) were excluded from analyses.

#### First-Level fMRI Models

Modeling was performed using SPM12 (version 6678) (https://www.fil.ion.ucl.ac.uk/spm)and custom MATLAB scripts. Temporal band-pass was set to the hemodynamic response function (HRF) and 128 s for low and high pass, respectively. Regressors were convolved with a canonical HRF and temporal derivative. Nuisance variates included framewise displacement, motion parameters, cerebrospinal fluid time-series, and instantaneous pulse and respiration. ICA-AROMA (Pruim et al., 2015) was used to extract other potential sources of noise (e.g., white matter signal) and these were also included as nuisance variates, consistent with recent recommendations (Bijsterbosch et al., 2020). Volumes with framewise displacement >0.5 mm were censored. ***Anxiety Provocation.*** A detailed description of the modeling procedure is available elsewhere (Hur et al., 2020b). In brief, the anxiety-provocation task was modeled using variable-duration rectangular regressors time-locked to the anticipation epochs of Threat or Safety trials, to the presentation of aversive or benign stimulation, and to rating trials (**Supplementary Figure S1**). Volumes coincident with aversive stimulation were censored. ***Threat-Related Faces.*** The faces task was modeled using fixed-duration rectangular regressors time-locked to the blocks of angry, fearful, or happy faces (**Supplementary Figure S2**).

#### Brain Metrics

To fuse the fMRI and EMA data-streams, regions of interest were functionally prescribed at the group level based on significant task effects in whole-brain voxel-wise analyses (FDR *q* < 0.05, whole-brain corrected; see below) within anatomical regions selected based on prior large-scale studies and meta-analyses (e.g., Chavanne & Robinson, 2021; Hur et al., 2018; Hur et al., 2020b), consistent with recent recommendations (Bijsterbosch et al., 2020). It merits comment that this is a conservative approach. This reflects the fact that task effects (Student’s t) are estimated by dividing the mean within-subjects difference (e.g., Threat vs. Safety) by the between-subjects variation. All else being equal, regions showing stronger task effects will tend to show less between-subject variance, constraining relations with external variables (e.g., momentary negative affect). For each region (e.g., MCC) and subject, regression coefficients were extracted and averaged across voxels using cubical masks centered on the local maxima identified in voxelwise analyses (**Figure 2**). A ‘faces-only’ ROI (7 voxels, 56 mm^3^) was used for subcortical regions, whereas a larger ‘faces, edges, and corners’ ROI (27 voxels, 216 mm^3^) was used for cortical regions, consistent with prior EMA-fMRI research (e.g., Lopez, Hofmann, Wagner, Kelley, & Heatherton, 2014). As a validity check, one-sample *t*-tests were used to confirm the sign and magnitude of ROI-extracted values (not reported). ***Anxiety Provocation.*** The goal of our study was to test the relevance of the threat-anticipation network identified in prior work to negative affect in the real world (Hur et al., 2020; Shackman & Fox, 2021). Accordingly, for the anxiety-provocation paradigm, analyses focused on activation during the anticipation of aversive compared to benign stimulation. Regression coefficients for the resulting ‘Threat-versus-Safety’ contrast were extracted and averaged across voxels for the dorsal amygdala, BST, PAG, MCC, AI, FrO, and dlPFC. With the exception of the PAG—a small midline structure—coefficients were separately extracted for the left and right hemispheres. To minimize the number of comparisons and maximize statistical power, a factor analysis with principal components extraction and oblimin rotation was used to reduce data dimensionality. Here the ROI-level data serve as manifest indicators of a *theory-driven* circuit-of-interest and factor analysis provides a principled means of distinguishing relevant sub-components or facets. Consistent with recent methodological recommendations (Lim & Jahng, 2019), parallel analysis (PA)—which compares the eigenvalues of the sample correlation matrix to those obtained using random uncorrelated standardized normal variables— was used to determine the number of factors to retain. PA was implemented using the *nFactors* package (version 2.4.1.) for *R*. Visual inspection of the scree plot, the Kaiser criterion (eigenvalue > 1), and PA all indicated the presence of 2 factors, with frontocortical ROIs loading on the first factor and subcortical ROIs loading on the second (**Supplementary Table S1**). Similar results were yielded using varimax rotation (not reported). Based on this, we standardized (*z*-transformation) and averaged the ROI values to create composite measures of anxiety-related frontocortical (α = .89) and subcortical activity (α = .82) for each subject. Although this approach is widely used in the social and biological sciences (e.g., Doré, Weber, & Ochsner, 2017; Fox et al., 2015; Lopez et al., 2017; Moffitt et al., 2011)—and had the advantage of reducing the neuroimaging dataset from tens of thousands of voxels to two regional composites—it merits comment that the observed pattern of factor loadings necessarily reflects a combination of biological and measurement differences (e.g., signal-to-noise) across regions. To clarify the unique contribution of specific regions (e.g., MCC), follow-up tests employed regional composites created by averaging the standardized ROI values for the left and right hemispheres. ***Threat-Related Faces.*** For the faces paradigm, analyses focused on dorsal amygdala reactivity to threat-related (i.e., fearful or angry) faces compared to places, consistent with prior work (e.g., Swartz, Knodt, Radtke, & Hariri, 2015). Mean coefficients were separately extracted for the left and right amygdalae, standardized, and averaged for each subject (α = .82).

**Figure 2.**
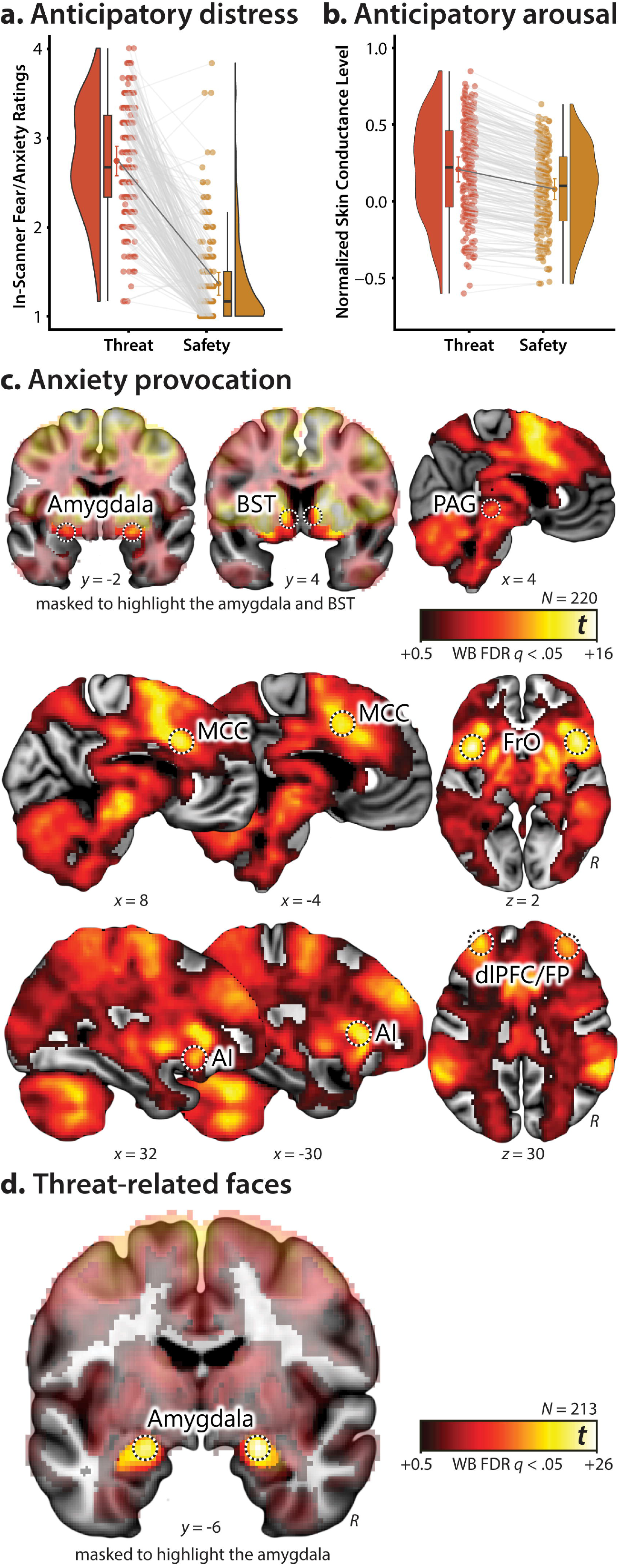
Laboratory measures for the anxiety-provocation (threat-anticipation) and threat-related faces paradigms. The anxiety-provocation paradigm produced robust symptoms (panel ***a***: inscanner ratings) and signs (panel ***b***: skin conductance) of anticipatory anxiety, confirming validity. Figures depict the data (dots and gray lines; *individual subjects*), density distribution (*bean plots*), Bayesian 95% highest density interval (HDI; *rectangular bands*), and mean (*bold black bars within the rectangular bands*) for each measure and condition. HDIs permit population-generalizable visual inferences about mean differences and were estimated using 1,000 samples from a posterior Gaussian distribution. The anxiety-provocation paradigm was also associated with increased activation (Threat > Safety; FDR *q*<.05, whole-brain corrected) in subcortical and frontocortical brain regions (panel c). The presentation of threat-related faces was associated with increased activation in the dorsal amygdala (panel ***d***: Threat-Related Faces > Places; FDR *q*<.05, whole-brain corrected). Note: The amygdala and BST panels are masked to highlight suprathreshold voxels in the relevant regions. Black-and-white rings indicate the regions used for fMRI-EMA analyses (see the Method and **Supplementary Tables S2-S3** for details). Abbreviations—AI, anterior insula; BST, bed nucleus of the stria terminalis; dlPFC, dorsolateral prefrontal cortex; FP, frontal pole; FrO, frontal operculum; MCC, midcingulate cortex; PAG, periaqueductal grey; R, right; WB, whole-brain corrected.

### Hypothesis Testing Strategy

Unless noted otherwise, hypothesis testing was performed using SPSS (version 24.0.0.0). To guard against error, a second analyst independently analyzed and confirmed key results.

#### In-Scanner Distress Ratings and Skin Conductance

To confirm the validity of the anxiety-provocation paradigm, repeated-measures GLMs were used to test differences between Threat and Safety anticipation. Unfortunately, the sparse nature of the in-scanner ratings protocol (i.e., two ratings per valence per scan) precluded meaningful analyses of concurrent brain-behavior relations. Raincloud plots were generated using open-source code (Allen, Poggiali, Whitaker, Marshall, & Kievit, 2019; van Langen, 2020).

#### fMRI

Standard whole-brain voxelwise GLMs with random effects were computed using SPM12 and used to assess activation to the anxiety-provocation (threat-anticipation) and emotional-faces paradigms. Significance was assessed using FDR *q*<.05, whole-brain corrected. Some figures were created using MRIcron (http://people.cas.sc.edu/rorden/mricron) and MRIcroGL (https://www.nitrc.org/projects/mricrogl). Clusters and local maxima were labeled using a combination of the Allen Institute, Harvard–Oxford, and Mai atlases (Desikan et al., 2006; Frazier et al., 2005; Hawrylycz et al., 2012; Mai, Majtanik, & Paxinos, 2015; Makris et al., 2006) and a recently established consensus nomenclature (ten Donkelaar, Tzourio-Mazoyer, & Mai, 2018). Frontal operculum subdivisions were labeled using the nomenclature of Amunts (Amunts et al., 2010). Amygdala subdivisions were labeled using the atlases and recently developed ROIs (Tillman et al., 2018; Tyszka & Pauli, 2016).

#### Brain-EMA Fusion

As shown schematically in **Figure 1**, a series of HLMs—often termed ‘multilevel’ or ‘linear mixed’ models—was used to test relations between individual differences in task-related brain activity and moment-by-moment levels of *tonic* (stressor-independent) and *reactive* (stressor-dependent) negative affect, separately for each composite brain metric. HLM naturally handles the nested dependency and variable number of longitudinal assessments provided by each subject, unlike traditional repeated-measures GLM approaches (Nezlek, 2012; Shiffman, Stone, & Hufford, 2008).

Here, the EMA-derived *Negative Affect* (continuous) and *Stressor* (binary) ‘time-series’ (i.e., Level-1 variables) were nested within subjects (see ‘EMA Survey and Data Reduction’ for details). Intercepts were free to vary across subjects. *Brain* metrics were derived in the manner described above, grandmean centered, and served as continuous Level-2 predictors.

For hypothesis testing purposes, a single HLM—incorporating terms for tonic *and* reactive negative affect—was implemented for each of the composite brain metrics. This has the advantage of providing estimates for each kind of affect (e.g., tonic) while controlling for the other (e.g., reactive). The following equations outline the basic structure of the HLM in standard notation (Raudenbush & Bryk, 2002). At the first level, *Negative Affect* during EMA *t* for individual *i* was modeled as a function of *Stressor* exposure, with the absence of *Stressor* exposure serving as the reference category (‘baseline’):

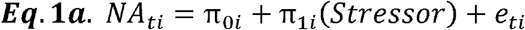

At the second level of the HLM, relations between *Stressors* and *Negative Affect* was modeled as a function of individual differences of the focal *Brain* metric:

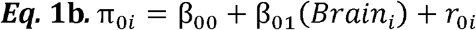

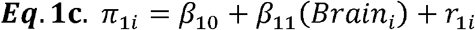

From a conceptual perspective, individual differences in *reactive Negative Affect* were estimated using a binary reference function (i.e., time-series) indicating the self-reported presence or absence of *Stressor* exposure at each EMA (Equation 1a). Individual differences in *tonic Negative Affect* were indexed by the intercept term in Equation 1a (i.e., each subject’s average mood, controlling for stressor-dependent fluctuations).

HLMs were computed using the SPSS default covariance structure (variance components) and restricted maximum likelihood estimates. Standardized (*z*-transformed) variables were used for all analyses. For illustrative purposes, significant interactions are depicted for extreme values (±1 *SD*) of the relevant brain metric (Cohen, Cohen, West, & Aiken, 2003). The Šidák procedure was used to determine corrected two-tailed significance thresholds for family set of tests that encompassed multiple brain metrics derived from a single task (e.g., subcortical and frontocortical composites; Šidák, 1967).

Using the same general approach, follow-up analyses were used to determine whether significant brain-EMA associations generalize to positive affect and positive events, which would suggest a broader functional role. To gauge the added explanatory value (‘incremental validity’) of significant brain metrics, we re-computed the relevant HLM after incorporating an alternative brain metric (e.g., amygdala reactivity to threat-related faces) or our multi-occasion index of trait negative emotionality, conceptually akin to performing a simultaneous multiple regression. Finally, to enhance interpretability, we decomposed significant brain-EMA associations by performing follow-up analyses for the anxious and depressed facets of negative affect, and for the brain regions (e.g., MCC) included in the relevant composite. Conclusions remained unchanged using square-root transformed negative affect or controlling for variation in EMA compliance (not reported).

## RESULTS

We first confirmed that the anxiety-provocation and threat-related faces paradigms had the intended effects on behavior and brain function. As expected, waiting to receive aversive stimulation was associated with robust increases in subjective symptoms of distress (in-scanner ratings of fear and anxiety) and objective signs of arousal (skin conductance), *t*s(219) > 27.00, *p*s < 0.001 (**Figures 2a-b**). As shown in **Figure 2c** and detailed in **Supplementary Table S2**, voxelwise GLMs focused on the period of threat anticipation revealed significantly increased activity in key *subcortical* (dorsal amygdala, BST, PAG) and *frontocortical* regions (MCC; FrO; AI; and dlPFC extending into the frontal pole, FP) of the threat-anticipation network (FDR q<.05, whole-brain corrected; Hur et al., 2020b; Shackman & Fox, 2021). Likewise, the presentation of threat-related faces was associated with increased activity in the dorsal amygdala (**Figure 2d**), fusiform gyrus, and other regions of the ventral visual cortex (**Supplementary Table S3**).

To fuse the fMRI and EMA data-streams, we first extracted measures of task-related activation (i.e., regression coefficients) from ROIs centered on peak voxels in key anatomical regions for each task and subject (*black-and-white rings* in **Figure 2**; see **Supplementary Tables S2-3** for coordinates). To reduce the number of comparisons, a factor analysis was used to guide the construction of composite measures for the anxiety-provocation task. Results revealed two latent factors, with subcortical regions (dorsal amygdala, BST, and PAG) loading on one factor and frontocortical regions (MCC, AI, FrO, and dlPFC/FP) loading on the other (**Supplementary Table S1**). Based on this, we standardized and averaged the ROI values to create composite measures of subcortical (α = .85) and frontocortical (α = .89) activation for each subject. A similar approach was used to create a bilateral amygdala composite for the threat-related faces task (α = .82). As shown schematically in **Figure 1**, a series of HLMs was then used to test relations between individual differences in task-related activation and momentary levels of *tonic* (stressor-independent) and *reactive* (stressor-dependent) negative affect. Unlike traditional repeated-measures GLM approaches, HLM naturally handles the nested dependency and variable number of EMAs contributed by each subject (Shiffman et al., 2008).

We first examined the anxiety-provocation task. As shown in **Figure 3a** and **Table 1**, subcortical and frontocortical activation during threat anticipation was unrelated to tonic (stressor-independent) levels of negative affect (*p*s > .64). Outside the laboratory, stressor exposure was associated with a momentary increase in negative affect (*p*s < .001). Although subcortical activation during the threat-anticipation task was unrelated to the magnitude of this stressor-dependent distress (*p* = .43), frontocortical reactivity *was* significantly related (*β* = −.07, *SE* = .03, *p* = .02), and remained significant after applying a principled correction for the number of ROIs examined (Šidák α_Critical_ = .025). Closer inspection indicated that individuals showing *more* activation in frontocortical regions when anticipating aversive stimulation experienced *lower* levels of negative affect in the moments following stressor exposure, consistent with a regulatory role (**Figure 3a**, ***inset***). This association remained significant (*β* = −.09, *p* = .03) after controlling for individual differences in subcortical reactivity, which were themselves not related to reactive distress (*p* = .43; **Supplementary Table S4**). Similar associations with frontocortical reactivity were evident for the anxious and depressed facets of momentary negative affect (*β*s = −.07, *p*s = .04-.05; **Supplementary Table S5**). Follow-up tests indicated that frontocortical reactivity was unrelated to the frequency of momentary stressors (*r* = −.01, *p* = .91), and relations between frontocortical activation and reactive distress remained significant after controlling for individual differences in the frequency of stressor exposure (*β* = −.07, *p* = .02; **Supplementary Table S6**).

**Figure 3.**
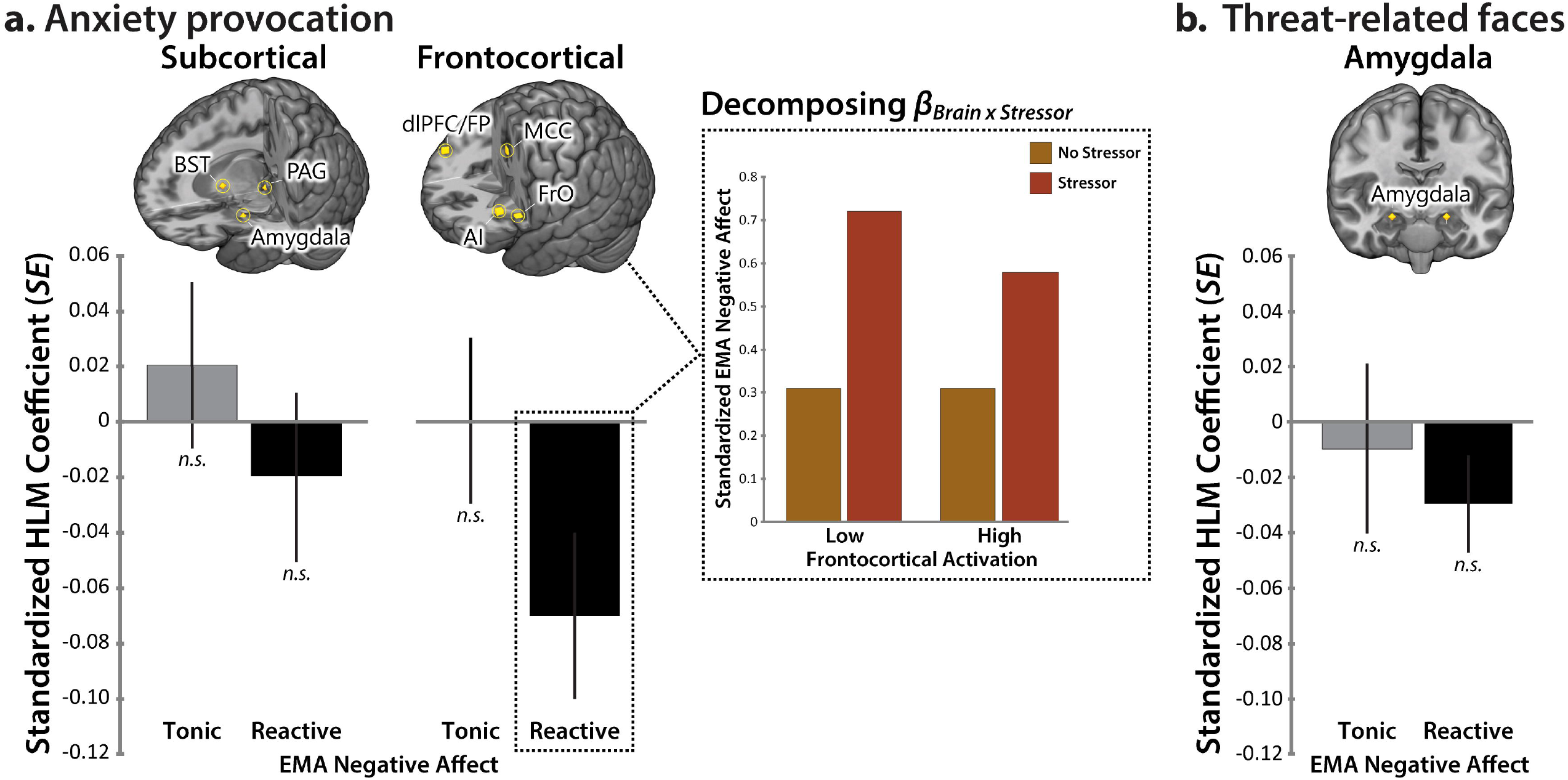
Frontocortical reactivity to the anxiety-provocation paradigm is associated with diminished stressor reactivity in the real world. ***a.*** EMA-fMRI fusion for the anxiety-provocation paradigm. Figure depicts standardized HLM coefficients for EMA-derived measures of tonic (stressor-independent; *gray*) and reactive (stressor-dependent; *black*) negative affect. The left side of the panel depicts non-significant results for the subcortical composite. The right side depicts significant results for the frontocortical composite. Error bars indicate the *SE*. Rendered brains depict the regions contributing to each composite (*gold*). Some regions are not visible. Inset depicts the decomposition of the Brain × Stressor interaction for the frontocortical composite. For illustrative purposes, HLM-predicted levels of negative affect are depicted for assessments associated with the absence (*orange*) or presence (*red*) of recent real-world stressors, separately for extreme levels (±1 *SD*) of frontocortical activation. Individuals with greater frontocortical reactivity to the anxiety-provocation (threat-anticipation) task in the laboratory experienced lower levels of negative affect in the moments during and following exposure to daily stressors. Hypothesis testing relied on continuous measures. ***b.*** EMA-fMRI fusion for the threat-related faces paradigm. Conventions are identical to panel *a*, and neither effect was significant. **Abbreviations**—AI, anterior insula; BST, bed nucleus of the stria terminalis; dlPFC, dorsolateral prefrontal cortex; FP, frontal pole; FrO, frontal operculum; MCC, midcingulate cortex; *n.s.*, not significant (*p* > .05); PAG, periaqueductal grey.

**Table 1.**
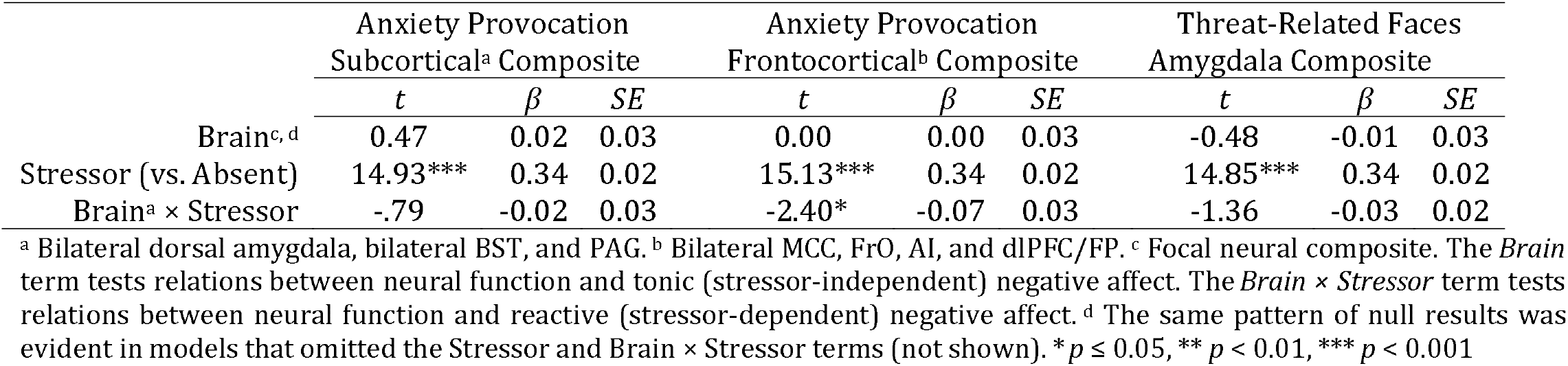
Relations between laboratory measures of brain function and real-world negative affect.

Next, we examined the emotional faces task. Variation in amygdala reactivity to threat-related faces in the laboratory was unrelated to either tonic (stressor-independent) or reactive (stressor-dependent) levels of negative affect in the field (*p*s > .17; **Figure 3c** and **Table 1**). Consistent with this null result, relations between frontocortical activation during the anxiety-provocation paradigm and reactive distress remained significant after controlling for amygdala reactivity to threat-related faces, underscoring the unique explanatory contribution of frontocortical function (*β* = −.07, *p* = .03; **Supplementary Table S7**).

These results demonstrate that heightened frontocortical activation while waiting for aversive stimulation is associated with lower levels of negative affect during and following exposure to everyday stressors. This is consistent with either a narrow role in dampening distress or a broader role in regulating emotion. To address this ambiguity, we used HLM to perform a parallel analysis for positive affect and positive events. Results indicated that frontocortical activation was unrelated to either tonic or reactive positive affect, consistent with a narrower regulatory function (*p*s > .14; **Supplementary Table S8**).

In head-to-head comparisons of criterion validity, simple paper-and-pencil measures often outperform more sophisticated brain imaging measures (Shackman & Fox, 2018). To clarify the explanatory value of frontocortical activation, we computed a new HLM that used a combination of frontocortical reactivity *and* a multi-scale, multi-occasion composite measure of trait negative emotionality (see Method) to explain momentary negative affect (conceptually akin to a multiple regression). As expected, negativity emotionality promoted distress; individuals with a more negative temperament experienced higher levels of tonic *and* reactive negative affect in their daily lives (*p*s < .001) (Bolger, 1990; Bolger & Schilling, 1991; Gross et al., 1998; Shackman et al., 2016; Thake & Zelenski, 2013). But more importantly, relations between frontocortical activation and reactive (stressor-dependent) negative affect remained significant after controlling for differences in negative emotionality (*β* = −.06, *p* = .03; **Supplementary Table S9**). In other words, frontocortical reactivity to the anxiety-provocation task accounted for variation in momentary distress above-and-beyond that explained by a traditional paper-and-pencil measure of emotional reactivity, underscoring the added explanatory value (‘incremental validity’) of the neuroimaging metric.

Multiregion composites have a number of psychometric and statistical advantages, but, by their nature, do not speak to the contributions of their constituent regions. To address this, we used a series of HLMs to decompose the frontocortical composite and determine the relevance of individual regions to momentary negative affect. As shown in **Figure 4** and detailed in **Supplementary Table S10**, individuals showing greater activation in either the MCC or FrO during the anxiety-provocation task experienced significantly lower levels of negative affect in the moments during and following stressor exposure in the field (*p*s < .01). Both regional associations remained significant after correcting for the number of frontocortical ROIs examined (Šidák α_Critical_ = .013). None of the other frontocortical brain-EMA associations was significant (*p*s > .11). Follow-up tests indicated that relations between anxiety-related MCC activation and reactive negative affect remained significant after controlling for variation in AI and dlPFC activation (*β* = −.08, *p* = .04; **Supplementary Table S11**). The same pattern was evident for the FrO (*β* = −.07, *p* = .05; **Supplementary Table S12**). AI and dlPFC were both unrelated to reactive negative affect in these models (*p*s > .56). Neither cingulo-opercular region was significantly related to reactive negative affect after controlling for variation in the other (e.g., MCC controlling for FrO; *p*s > .28; **Supplementary Table S13**), consistent with the substantial association between anxiety-related MCC and FrO activation (*r* = .72, *p* < .001). Taken together, these observations suggest that, among the frontocortical regions recruited by the anxiety-provocation paradigm, cingulo-opercular activation is most closely related to real-world variation in reactive (stressor-dependent) negative affect.

**Figure 4.**
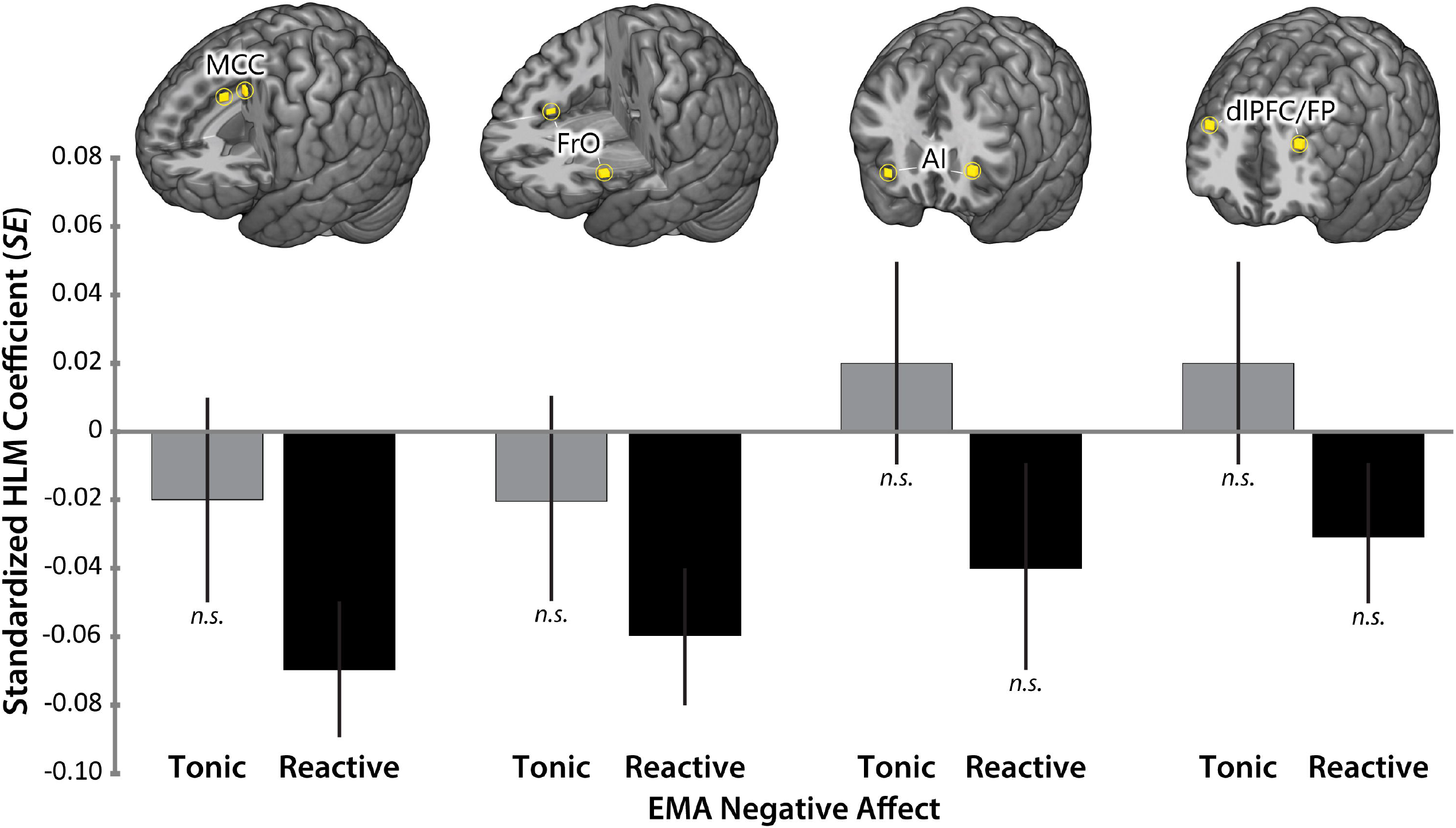
MCC and FrO reactivity to the anxiety-provocation paradigm is associated with diminished stressor reactivity in the real world. Conventions are identical to **Figure 3a**. ***a.*** MCC. ***b.*** FrO. ***c.*** AI. ***d.*** dlPFC/FP. **Abbreviations**—AI, anterior insula; dlPFC, dorsolateral prefrontal cortex; FP, frontal pole; FrO, frontal operculum; MCC, midcingulate cortex; *n.s.*, not significant (*p* > .05).

Our findings raise the possibility that the MCC and FrO represent a meaningful functional circuit. Consistent with this possibility, a series of supplementary analyses showed that these regions show robust coupling (‘intrinsic functional connectivity’) at rest *and* are consistently co-activated across a range of experimental challenges (Laird et al., 2013; Yeo et al., 2011). These findings are described in more detail in the Supplement and visually summarized in **Supplementary Figure S3**.

## DISCUSSION

Anxiety, sadness, and other negative feelings are a hallmark of the human condition and play a central role in contemporary theories of decision making, development, interpersonal processes, personality, psychopathology, and well-being (Fox et al., 2018). Recent work has begun to reveal the neural systems governing the expression and regulation of negative affect, but the relevance of these tantalizing laboratory discoveries to the real world has remained uncertain. Here, we used a combination of fMRI and EMA data to demonstrate that individuals showing greater frontocortical activation during a well-established anxiety-provocation (threat-anticipation) task experience dampened reactive (stressor-dependent) distress in their daily lives (**Figure 3**). Frontocortical activation was not significantly related to momentary positive affect or to tonic (stressor-independent) negative affect, suggesting a relatively narrow functional role. In a simultaneous HLM, frontocortical activation accounted for variation in daily distress above-and-beyond a conventional psychometric measure of trait negative emotionality, underscoring its added explanatory value. Follow-up analyses indicated that, among the frontocortical regions that we examined in detail, this association predominantly reflected heightened engagement of a functionally coherent cingulo-opercular (MCC and FrO) circuit (**Figure 4** and **Supplementary Figure S3**).

These findings have implications for understanding the neural systems underlying negative affect. There is ample evidence that the MCC and FrO are recruited by distress-eliciting laboratory challenges, including instructed threat-of-shock, Pavlovian threat conditioning, and physical pain (Chavanne & Robinson, 2021; Fullana et al., 2016; Shackman et al., 2011; Xu et al., 2020). This has led some to conclude that the cingulo-opercular network plays a role in assembling and expressing negative affect (Etkin, Buchel, & Gross, 2015; Hinojosa, Kaur, VanElzakker, & Shin, 2019; Milad & Quirk, 2012). Yet recent metaanalyses make it clear that the MCC and FrO are also recruited by tasks that demand controlled processing and behavioral flexibility, including popular assays of cognitive conflict (e.g., go/no-go) and emotion regulation (Langner et al., 2018; Morawetz et al., 2020; Shackman et al., 2011; Uddin, *in press*). Furthermore, MCC activation tracks variation in both the cognitive demands associated with deliberate emotion regulation *and* the degree of regulatory success (Urry, van Reekum, Johnstone, & Davidson, 2009). These observations suggest that cingulo-opercular activation during aversive laboratory challenges reflects spontaneous efforts to down-regulate or inhibit distress, a process that some have termed ‘implicit’ emotion regulation (Shackman & Lapate, 2018; Van Reekum & Johnstone, 2018). Our results—which demonstrate that heightened cingulo-opercular reactivity to an anxiety provocation task is associated with lower levels of reactive (stressor-dependent) distress in daily life—are consistent with this hypothesis. While mechanistic evidence is scant, the present findings are well aligned with work showing that surgical damage to the MCC (‘cingulotomy’) is associated with increased emotional reactivity to painful stimuli in humans (Davis, Hutchison, Lozano, & Dostrovsky, 1994; Greenspan et al., 2008). Likewise, focal inactivation of the posterior MCC transiently increases defensive responses to intruder and snake threats in monkeys (Rahman et al., *in press*). Together, this body of data is consistent with conceptual models that emphasize the importance of the cingulo-opercular network for flexibly controlling cognition, emotion, and action in situations where automatic or habitual responses are inadequate, as when there is competition between plausible alternative actions or between action and inaction (e.g., passively respond to emotional challenges vs. deliberately regulate the response) (Shackman et al., 2011; Uddin, *in press*). The present results help to extend this framework from the artificial confines of the neuroimaging laboratory to the real world.

Clearly important challenges remain. First, the present study was focused on understanding the relevance of anxiety-related brain function to momentary levels of negative affect in the daily lives of young adults. Moving forward, it will be useful to expand this to encompass nationally representative samples (LeWinn, Sheridan, Keyes, Hamilton, & McLaughlin, 2017) and concurrent relations with trial-by-trial fluctuations in negative affect, an analytic approach not permitted by the in-scanner ratings used here (Geuter et al., 2020; Lim, Padmala, & Pessoa, 2009). Second, our results indicate that subcortical reactivity to threat-anticipation and amygdala reactivity to threat-related faces in the laboratory are unrelated to distress in the field, despite a relatively well-powered sample. While there are a number of possible explanations, this null effect is not unprecedented. Three recent large-sample studies (Duke Neurogenetics Study: *n* = 1,256; HCP: *n* = 319; Minnesota Twin Study: *n* = 548) failed to detect credible relations between amygdala reactivity to threat-related faces and individual differences in negative emotionality (MacDuffie, Knodt, Radtke, Strauman, & Hariri, 2019; Silverman et al., 2019; West, Burgess, Dust, Kandala, & Barch, *in press*). Does this mean that the amygdala, BST, and PAG are unrelated to negative affect? No, mechanistic work in humans, monkeys, and rodents makes it abundantly clear that they are (Fox & Shackman, 2019; Hur et al., 2019). Instead, this work raises the possibility that conventional fMRI measures of emotion perception (*viewing photographs of fearful or angry faces*) and generation (*briefly waiting for aversive stimulation*) are suboptimal probes of the aspects of subcortical function most relevant to everyday affect (i.e., ‘wrong’ assay) (Puccetti et al., 2021; Sicorello et al., 2021). Alternatively, it could be that isolated regional measures of subcortical function are only weakly predictive of conscious feelings of negative affect and, hence, to typical state, trait, and clinical assessments (Brown, Lau, & LeDoux, 2019; Chang et al., 2015; LeDoux & Pine, 2016; Shackman & Fox, 2018). Adjudicating between these possibilities is a key challenge for future research.

Anxiety disorders and depression are defined, diagnosed, and treated on the basis of negative feelings experienced in the midst of daily life. These disorders impose a staggering burden on global public health and existing treatments are inconsistently effective, underscoring the urgency of developing a deeper understanding of the underlying neurobiology (Dieleman et al., 2020; Ormel, Kessler, & Schoevers, 2019; Sartori & Singewald, 2019; Vos et al., 2020). The present findings highlight the relevance of cingulo-opercular function for real-world distress. These observations lay the groundwork for the kinds of prospective-longitudinal and mechanistic studies that will be necessary to determine causation and develop more effective interventions. A relatively large sample and best-practice approaches to data acquisition, processing, and analysis enhance confidence in the robustness and translational relevance of these results.

## Supporting information

Supplementary Method and Results

## ACKNOWLEDGEMENTS

This work was supported by the NIH (MH107444, MH121409, MH121735). Authors declare no conflicts of interest.

## RESOURCE SHARING

Raw data (https://nda.nih.gov) and processed EMA data (https://github.com/dr-consulting/shackman-umd-pax-ema-pub) are available. Key neuroimaging maps have been or will be uploaded to NeuroVault.org.

