## Supplementary Method and Results for "Anxiety-related frontocortical activity is associated with dampened stressor reactivity in the real world"

Juyoen Hur^1^*

Manuel Kuhn^2^*

Shannon E. Grogans^3^*

Allegra S. Anderson^6^

Samiha Islam^7^

Hyung Cho Kim^3,4^

Rachael M. Tillman^3^

Andrew S. Fox^8,9^

Jason F. Smith^3^†

Kathryn A. DeYoung^3^†

Alexander J. Shackman^3,4,5^†

^1^Department of Psychology, Yonsei University, Seoul 03722, Republic of Korea. ^2^Center for Depression, Anxiety and Stress Research, McLean Hospital, Harvard Medical School, Belmont, MA 02478 USA. Department of ^3^Psychology; ^4^Neuroscience and Cognitive Science Program; and ^5^Maryland Neuroimaging Center, University of Maryland, College Park, MD 20742 USA. ^6^Department of Psychological Sciences, Vanderbilt University, Nashville, TN 37240 USA. ^7^Department of Psychology, University of Pennsylvania, Philadelphia, PA USA. ^8^Department of Psychology and ^9^California National Primate Research Center, University of California, Davis, CA 95616 USA

* contributed equally * | † contributed equally

**Address Correspondence to:**

Juyoen Hur or Alexander J. Shackman

**Table of Contents**

**Supplementary Details**

Detailed Acknowledgements	p. 3

Detailed Author Contributions	p. 3

**Brain Imaging Paradigms**

Figure S1. Anxiety-Provocation (Threat-Anticipation) Paradigm	p. 4

Figure S2. Threat-Related Faces Paradigm	p. 5

**Dimensionality Reduction for Anxiety-Provocation (Threat-Anticipation) fMRI Data**

Table S1. Oblimin-rotated factor loadings for anxiety-provocation (threat-anticipation) regions of interest	p. 6

**Brain Imaging Results**

Table S2. Threat compared to Safety anticipation, cluster descriptive statistics	p. 6

Table S3. Threat-related faces compared to places, cluster descriptive statistics	p. 8

**Brain-EMA Results**

Table S4. Frontocortical reactivity to threat and negative affect, controlling for subcortical reactivity	p. 10

Table S5. Frontocortical reactivity to threat and anxiety and depression (facets of negative affect)	p. 11

Table S6. Frontocortical reactivity to threat and negative affect, controlling for stressor frequency	p. 11

Table S7. Frontocortical reactivity to threat and negative affect, controlling for amygdala reactivity to faces	p. 11

Table S8. Frontocortical reactivity to threat and positive affect	p. 12

Table S9. Frontocortical reactivity to threat and negative affect, controlling for trait negative emotionality	p. 12

Table S10. Frontocortical regional reactivity to threat and negative affect	p. 12

Table S11. MCC reactivity to threat and negative affect, controlling for AI and dlPFC/FP reactivity	p. 13

Table S12. FrO reactivity to threat and negative affect, controlling for AI and dlPFC/FP reactivity	p. 13

Table S13. MCC and FrO reactivity to threat and negative affect, controlling for the other region	p. 13

**Supplementary Method for Meta-Analytic Analyses of the Cingulo-Opercular Circuit**	p. 14

**Meta-Analytic Results and Discussion**	p. 14-16

Figure S3. The MCC and FrO form a coherent functional neuroanatomical circuit	p. 16

**Supplementary References**	p. 17

**Detailed Acknowledgements**

Authors acknowledge assistance and critical feedback from A. Antonacci, M. Barstead, L. Friedman, J. Furcolo, M. Gamer, C. Grubb, R. Hum, C. Kaplan, T. Kashdan, J. Kuang, C. Lejuez, D. Limon, B. Nacewicz, L. Pessoa, S. Rose, J. Swayambunathan, A. Vogel, J. Wedlock, members of the Affective and Translational Neuroscience laboratory, the staff of the Maryland Neuroimaging Center, and the Office of the Registrar at the University of Maryland. This work was supported by the California National Primate Center; National Institute of Mental Health (MH107444, MH121409, and MH121735); University of California, Davis; and University of Maryland, College Park. Authors declare no conflicts of interest.

**Detailed Author Contributions**

A.J.S., K.A.D., and J.F.S. designed the overall study. J.F.S. and A.J.S. developed and optimized the imaging paradigm. K.A.D. and A.J.S. developed and optimized the EMA paradigm. K.A.D. managed data collection and study administration. K.A.D., J.F.S., A.S.A, S.I., and R.M.T. collected data. K.A.D. and J.H. processed and analyzed EMA data. J.F.S. and M.K. developed data processing and analytic software for imaging analyses. J.H., J.F.S., H.C.K., and R.M.T. processed imaging data. J.H., M.K., J.F.S., and A.J.S. developed the strategy for the imaging analyses. M.K., J.H., and J.F.S. analyzed imaging data. J.H. and A.J.S. developed the strategy for the EMA-imaging analyses. J.H., M.K., A.S.F., S.E.G., and A.J.S. interpreted data. S.E.G., J.H., A.S.F., and A.J.S. wrote the paper. S.E.G., M.K., J.H., and A.J.S. created figures. M.K., J.H., and A.J.S. created tables. A.J.S. funded and supervised all aspects of the study. All authors contributed to reviewing and revising the paper and approved the final version.

***Continued…***

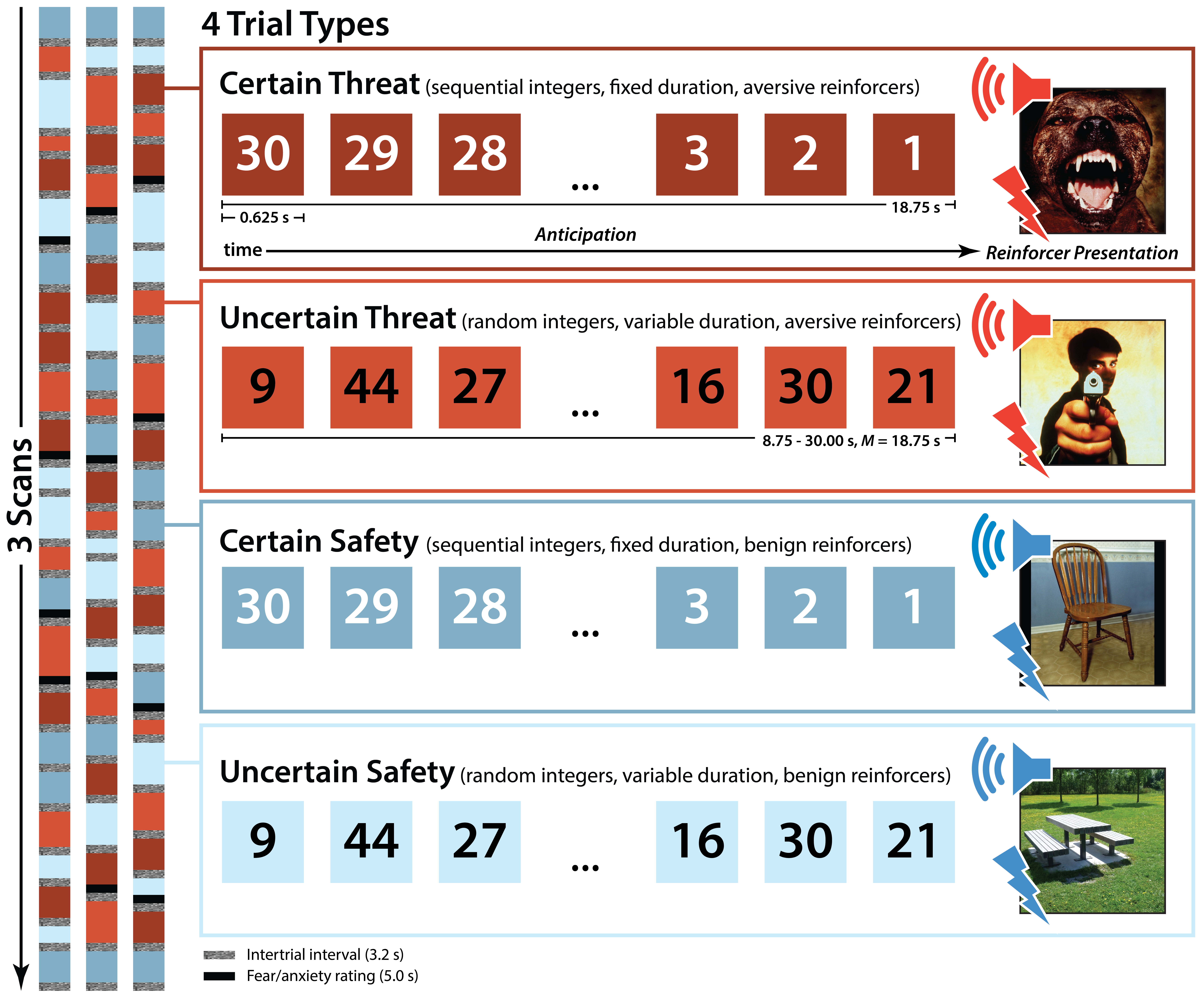

**Supplementary Figure S1**. **Anxiety-Provocation Paradigm.** The anxiety-provocation (threat-anticipation) task took the form of a 2 (*Valence:* Threat/Safety) × 2 (*Temporal Certainty:* Certain/Uncertain) randomized event-related design, as described in detail elsewhere (Hur et al., 2020). Subjects were completely informed about the task design and contingencies prior to scanning. The task was administered in 3 scans, with short breaks between scans. On Certain-Threat trials, subjects saw a descending stream of integers (‘count-down;’ e.g. 30, 29, 28...3, 2, 1) for 18.75 s. To ensure robust anxiety, this anticipation epoch always culminated with the delivery of a noxious electric shock, unpleasant photographic image (e.g., mutilated body), and thematically related audio clip (e.g., scream, gunshot). Uncertain-Threat trials were similar, but the integer stream was randomized and presented for an uncertain and variable duration (8.75-30.00 s; *M*=18.75 s). Subjects knew that something aversive was going to occur, but they had no way of knowing precisely *when*. The mean duration of the anticipation epoch was chosen to enhance detection of task-related differences in the blood oxygen level-dependent (BOLD) signal (Henson, 2007a), and was identical across trial types, ensuring an equal number of measurements. Safety trials were similar, but terminated with the delivery of benign reinforcers (e.g., just-perceptible electrical stimulation and neutral audiovisual stimuli). Valence was continuously signaled during the anticipation epoch by the background color of the display. White-noise visual masks (3.2 s) were presented between trials to minimize persistence of visual reinforcers in iconic memory. Subjects were periodically prompted to rate the intensity of negative affect (‘fear/anxiety’) experienced a few seconds earlier, during the *anticipation* period of the prior trial. Each condition was rated twice per scan. Skin conductance was continuously acquired. ***Design Consideration.*** The major aim of the present study was to understand the relevance of anxiety-related brain function to negative affect as it is experienced in the midst of everyday life. Prior work using this paradigm demonstrates that both kinds of threat anticipation—certain and uncertain—produce similar increases in anxious distress and arousal and recruit highly overlapping brain circuits (Hur et al., 2020). Therefore, the present analyses focused on the overall effect of threat anticipation—aggregated across the certainty manipulation—as detailed in the main report.

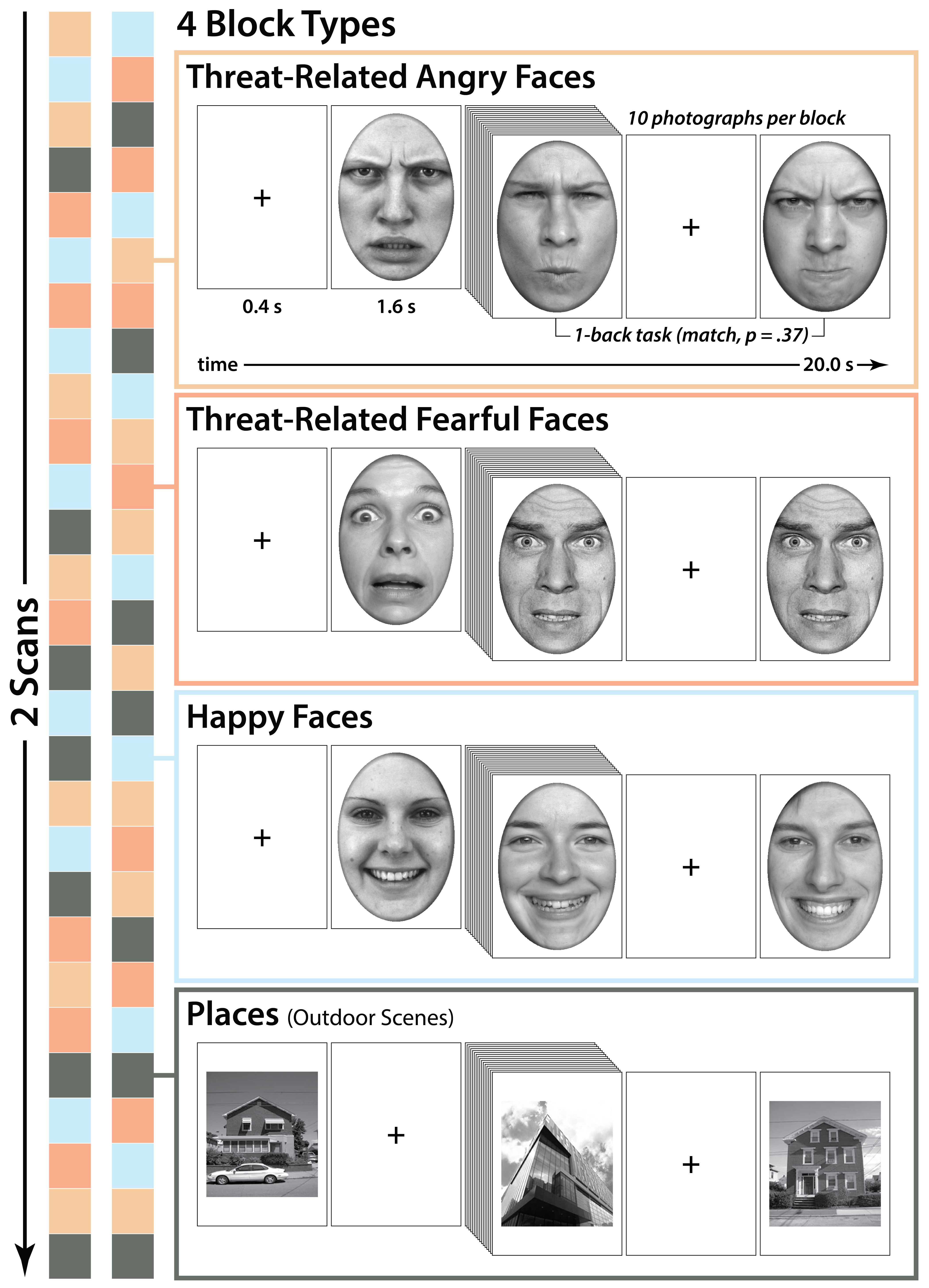

**Supplementary Figure S2**. **Emotional Faces Paradigm*.*** The faces task took the form of a pseudo-randomized block design and was administered in 2 scans, with a short break between scans. During each scan, subjects viewed photographs of adult models (half female) depicting angry faces, fearful faces, happy faces, or places (7 blocks/condition/scan). To maximize signal strength and homogeneity, and minimize potential habituation (Henson, 2007b; Maus, van Breukelen, Goebel, & Berger, 2010; Plichta et al., 2014), blocks consisted of 10 photographs (1.6 s) separated by fixation crosses (0.4 s). To ensure engagement, subjects judged whether the current photograph matched that presented on the prior trial (i.e., a ‘1-back’ continuous performance task). Matches occurred 37.1% of the time. To minimize potential habituation, each photograph was presented once or twice. Face stimuli were adapted from prior work (Gamer, Schmitz, Tittgemeyer, & Schilbach, 2013; Scheller, Büchel, & Gamer, 2012) and included photographs of prototypical emotional expressions from Ekman and Friesen’s Pictures of Facial Affect (Ekman & Friesen, 1976), the FACES database (Ebner, Riediger, & Lindenberger, 2010), the Karolinska Directed Emotional Faces database (<http://www.emotionlab.se/resources/kdef>), and the NimStim Face Stimulus Set (<https://www.macbrain.org/resources.htm>). Color photographs were converted to grayscale, brightness normalized, and masked to occlude non-facial features. Place stimuli consisted of grayscale photographs of outdoor scenes focused on single-family residential buildings (‘houses’) or urban commercial buildings (‘skyscrapers’), and were also adapted from prior work (Choi, Padmala, & Pessoa, 2012, 2015).

**Table S1. Oblimin-rotated factor loadings for anxiety-provocation (threat-anticipation) regions of interest.**

| **Region of Interest** | **Frontocortical Factor** | **Subcortical Factor** |
| --- | --- | --- |
| R Dorsolateral Prefrontal Cortex / Frontal Pole | **0.88** | -0.22 |
| L Frontal Operculum | **0.82** | 0.05 |
| L Midcingulate Cortex | **0.80** | 0.05 |
| L Dorsolateral Prefrontal Cortex / Frontal Pole | **0.77** | 0.01 |
| L Anterior Insula | **0.64** | 0.12 |
| R Midcingulate Cortex | **0.64** | 0.21 |
| R Frontal Operculum | **0.62** | 0.08 |
| R Anterior Insula | **0.42** | 0.35 |
| R Bed Nucleus of the Stria Terminalis | -0.02 | **0.83** |
| R Amygdala | -0.03 | **0.81** |
| L Bed Nucleus of the Stria Terminalis | 0.08 | **0.77** |
| L Amygdala | 0.02 | **0.76** |
| Periaqueductal Gray | 0.31 | **0.42** |

**Table S2. Descriptive statistics for clusters and local maxima showing greater activity during the anticipation of Threat compared to Safety (FDR *q*<.05, whole-brain corrected).**

| **mm^3^** | **Region** | ***t*** | ***x*** | ***y*** | ***z*** |
| --- | --- | --- | --- | --- | --- |
| 1,328,072 | L Dorsolateral Prefrontal Cortex/Frontal Pole^a^ | 15.05 | -30 | 50 | 30 |
|  | R Dorsolateral Prefrontal Cortex/Frontal Pole^a^ | 13.40 | 34 | 48 | 30 |
|  | L Middle Frontal Gyrus | 13.01 | -36 | 34 | 36 |
|  | R Middle Frontal Gyrus | 3.79 | 48 | 22 | 44 |
|  | L Frontal Orbital Cortex (Basal Operculum) | 16.11 | -34 | 28 | -4 |
|  | R Frontal Orbital Cortex (Basal Operculum) | 16.60 | 36 | 28 | 0 |
|  | L Anterior Insula (Anterior Insular Point)^a^ | 16.71 | -30 | 26 | 4 |
|  | R Anterior Insula (Orbitoinsularis Cortex)^a^ | 13.40 | 32 | 22 | -10 |
|  | R Midcingulate Cortex (Cingulate Gyrus, Cingulate Sulcus)^a^ | 17.08 | 8 | 20 | 34 |
|  | L Midcingulate Cortex (Paracingulate Gyrus)^a^ | 16.47 | -4 | 8 | 42 |
|  | R Frontal Operculum (Op 8)^a^ | 18.23 | 46 | 18 | 2 |
|  | L Frontal Operculum (Op 8)^a^ | 19.43 | -40 | 14 | 2 |
|  | R Inferior Frontal Gyrus, pars opercularis | 9.12 | 38 | 14 | 24 |
|  | R Caudate | 14.31 | 18 | 12 | 12 |
|  | R Putamen | 17.57 | 24 | 10 | 0 |
|  | L Putamen | 15.42 | -22 | 8 | 0 |
|  | L Temporal Pole | 5.79 | -40 | 10 | -38 |
|  | R Temporal Pole | 9.71 | 44 | 6 | -38 |
|  | L Bed Nucleus of Stria Terminalis^a,b^ | 9.61 | -8 | 4 | 0 |
|  | R Bed Nucleus of Stria Terminalis^a,b^ | 7.28 | 8 | 4 | 2 |
|  | R Superior Temporal Gyrus, anterior division | 3.00 | 54 | 4 | -14 |
|  | L Superior Frontal Gyrus | 15.81 | -14 | 2 | 68 |
|  | R Juxtapositional Lobule | 17.44 | 4 | 6 | 54 |
|  | L Juxtapositional Lobule | 15.39 | -6 | 2 | 52 |
|  | L Inferior Temporal Gyrus, anterior division | 6.30 | -50 | 0 | -34 |
|  | R Precentral Gyrus | 16.31 | 46 | 0 | 50 |
|  | L Amygdala^a,c^ (Medial Nucleus) | 5.93 | -22 | -2 | -12 |
|  | L Caudate | 14.15 | -14 | -2 | 16 |
|  | R Superior Frontal Gyrus | 16.67 | 12 | -2 | 72 |
|  | R Amygdala^a,d^ (Central, Medial Nuclei) | 8.03 | 22 | -2 | -12 |
|  | L Planum Polare | 4.76 | -42 | -4 | -18 |
|  | L Parahippocampal Gyrus, anterior division | 6.82 | -26 | -4 | -32 |
|  | L Precentral Gyrus | 13.42 | -34 | -6 | 60 |
|  | R Parahippocampal Gyrus, anterior division | 4.47 | 24 | -6 | -32 |
|  | L Pallidum | 12.81 | -18 | -10 | 0 |
|  | R Thalamus | 13.10 | 16 | -10 | 6 |
|  | L Thalamus | 8.50 | -6 | -20 | 16 |
|  | R Planum Polare | 5.77 | 42 | -10 | -12 |
|  | R Pallidum | 11.98 | 26 | -12 | 2 |
|  | L Temporal Fusiform Cortex, posterior | 2.74 | -32 | -14 | -42 |
|  | R Temporal Fusiform Cortex, posterior | 2.53 | 42 | -28 | -28 |
|  | R Posterior Cingulate Cortex | 12.35 | 8 | -20 | 40 |
|  | L Posterior Cingulate Cortex | 13.23 | -10 | -24 | 38 |
|  | L Brain Stem | 12.87 | -10 | -24 | -18 |
|  | R Brain Stem | 14.18 | 8 | -30 | -14 |
|  | R Parietal Operculum | 11.95 | 54 | -24 | 22 |
|  | L Inferior Temporal Gyrus, posterior | 4.03 | -60 | -28 | -22 |
|  | R Inferior Temporal Gyrus, posterior | 3.70 | 52 | -32 | -28 |
|  | R Postcentral Gyrus | 10.93 | 46 | -28 | 48 |
|  | R Periaqueductal Gray^a^ | 10.23 | 4 | -30 | -6 |
|  | R Middle Temporal Gyrus, posterior | 13.40 | 50 | -30 | -6 |
|  | L Middle Temporal Gyrus, posterior | 6.91 | -60 | -32 | -8 |
|  | R Hippocampus, posterior | 7.39 | 34 | -32 | -4 |
|  | L Postcentral Gyrus | 10.21 | -42 | -34 | 46 |
|  | R Supramarginal Gyrus, posterior | 16.44 | 60 | -42 | 34 |
|  | L Supramarginal Gyrus, posterior | 16.34 | -58 | -48 | 34 |
|  | L Inferior Temporal Gyrus, temporooccipital | 3.71 | -60 | -46 | -20 |
|  | R Inferior Temporal Gyrus, temporooccipital | 6.63 | 54 | -60 | -14 |
|  | R Superior Parietal Lobule | 13.07 | 24 | -48 | 66 |
|  | L Angular Gyrus | 9.98 | -50 | -54 | 50 |
|  | L Superior Parietal Lobule | 10.7 | -34 | -54 | 62 |
|  | L Precuneus | 10.14 | -10 | -56 | 60 |
|  | R Precuneus Cortex | 10.20 | 10 | -58 | 60 |
|  | R Middle Temporal Gyrus, temporooccipital | 9.67 | 44 | -56 | 8 |
|  | L Middle Temporal Gyrus, temporooccipital | 9.02 | -46 | -60 | 10 |
|  | L Lateral Occipital Cortex, superior | 10.27 | -32 | -60 | 60 |
|  | R Lateral Occipital Cortex, superior | 9.08 | 28 | -62 | 60 |
|  | R Lateral Occipital Cortex, inferior | 8.36 | 48 | -70 | -12 |
|  | L Lateral Occipital Cortex, inferior | 5.11 | -44 | -84 | -4 |
|  | L Lingual Gyrus | 7.18 | -12 | -86 | -12 |
|  | R Occipital Fusiform Gyrus | 4.69 | 22 | -88 | -6 |
|  | L Occipital Pole | 6.56 | -18 | -90 | 26 |
|  | R Occipital Pole | 8.54 | 28 | -90 | 10 |
| 56 | R Frontal Pole | 2.20 | 10 | 74 | 8 |
| 8 | R Inferior Temporal Gyrus, posterior | 1.87 | 54 | -18 | -30 |

^a^ Region used in EMA analyses. ^b^ Overlaps the BST sub-region anatomically defined by Theiss and colleagues (Theiss, Ridgewell, McHugo, Heckers, & Blackford, 2017). ^c^ Harvard-Oxford left amygdala, *p*=36%. ^c^ Harvard-Oxford right amygdala, *p*=56%.

**Table S3. Descriptive statistics for clusters and local maxima showing greater activity during threat-related faces compared to places (FDR *q*<.05, whole-brain corrected).**

| **mm^3^** | **Region** | ***t*** | ***x*** | ***y*** | ***z*** |
| --- | --- | --- | --- | --- | --- |
| 1,440,200 | R Frontal Pole | 11.42 | 4 | 58 | 24 |
|  | L Frontal Pole | 8.8 | -34 | 52 | 26 |
|  | R Frontal Medial Cortex | 13.57 | 4 | 54 | -14 |
|  | L Frontal Medial Cortex | 12.4 | -4 | 50 | -16 |
|  | R Inferior Frontal Gyrus, pars triangularis | 11.6 | 56 | 34 | 6 |
|  | L Inferior Frontal Gyrus, pars triangularis | 6.04 | -54 | 30 | 4 |
|  | R Frontal Operculum | 9.81 | 46 | 26 | 0 |
|  | R Middle Frontal Gyrus | 6.15 | 46 | 26 | 44 |
|  | L Middle Frontal Gyrus | 16.75 | -34 | -4 | 60 |
|  | R Pregenual Anterior Cingulate Cortex | 9.24 | 10 | 46 | -4 |
|  | L Midcingulate Cortex (Paracingulate Gyrus) | 12.64 | -2 | 20 | 38 |
|  | L Midcingulate Cortex (Cingulate Sulcus) | 11.59 | -6 | 22 | 30 |
|  | R Midcingulate Cortex (Cingulate Gyrus) | 12.17 | 6 | 16 | 34 |
|  | R Accumbens | 7.58 | 12 | 20 | -6 |
|  | L Accumbens | 7.42 | -10 | 16 | -6 |
|  | R Inferior Frontal Gyrus, pars opercularis | 11.57 | 46 | 16 | 28 |
|  | L Inferior Frontal Gyrus, pars opercularis | 9.03 | -46 | 16 | 26 |
|  | R Frontal Orbital Cortex (Gustatory Cortex) | 12.26 | 26 | 14 | -20 |
|  | L Frontal Orbital Cortex / Insula | 11.9 | -28 | 14 | -16 |
|  | L Superior Frontal Gyrus | 10.75 | -12 | 10 | 66 |
|  | R Temporal Pole | 10.4 | 36 | 4 | -40 |
|  | L Temporal Pole | 13.6 | -34 | 4 | -20 |
|  | R Middle Temporal Gyrus, anterior | 5.79 | 50 | 2 | -30 |
|  | L Middle Temporal Gyrus, anterior | 5.73 | -62 | 0 | -18 |
|  | R Insula | 16.44 | 38 | 2 | -18 |
|  | L Insula | 14.11 | -38 | 0 | -16 |
|  | L Superior Temporal Gyrus, anterior | 5.48 | -52 | 2 | -16 |
|  | R Precentral Gyrus | 18.73 | 46 | -2 | 50 |
|  | R Temporal Fusiform Cortex, anterior | 10.55 | 32 | -2 | -38 |
|  | L Pallidum | 7.19 | -14 | -2 | -2 |
|  | L Putamen | 10.22 | -24 | -2 | 4 |
|  | R Putamen | 12.18 | 30 | -22 | 2 |
|  | L Parahippocampal Gyrus, anterior | 10.76 | -30 | -2 | -34 |
|  | L Juxtapositional Lobule | 19.21 | -2 | -8 | 68 |
|  | R Juxtapositional Lobule | 17.24 | 0 | -4 | 60 |
|  | R Amygdala^a,b^ (Medial Nucleus) | 25.76 | 22 | -6 | -14 |
|  | L Amygdala^a,c^ (Medial Nucleus) | 22.78 | -20 | -6 | -14 |
|  | R Caudate | 12.05 | 16 | -6 | 20 |
|  | L Central Operculum | 9.82 | -40 | -6 | 14 |
|  | L Caudate | 12.8 | -16 | -8 | 20 |
|  | R Superior Frontal Gyrus | 16.69 | 20 | -10 | 72 |
|  | L Thalamus | 14.65 | -2 | -12 | 6 |
|  | R Thalamus | 16.07 | 6 | -16 | 8 |
|  | L Precentral Gyrus | 20.3 | -44 | -12 | 54 |
|  | R Middle Temporal Gyrus, posterior | 13.37 | 48 | -18 | -10 |
|  | R Posterior Cingulate Cortex | 13.35 | 2 | -18 | 32 |
|  | L Posterior Cingulate Cortex | 13.71 | -10 | -22 | 42 |
|  | R Planum Polare | 4.44 | 40 | -20 | -2 |
|  | R Inferior Temporal Gyrus, posterior | 4 | 48 | -22 | -26 |
|  | L Inferior Temporal Gyrus, posterior | 2.18 | -62 | -42 | -24 |
|  | L Superior Temporal Gyrus, posterior | 8.83 | -54 | -22 | -4 |
|  | R Supramarginal Gyrus, anterior | 12.56 | 52 | -24 | 40 |
|  | L Postcentral Gyrus | 20.04 | -42 | -24 | 54 |
|  | R Postcentral Gyrus | 16.84 | 12 | -32 | 72 |
|  | L Hippocampus | 5.92 | -26 | -28 | -12 |
|  | R Hippocampus | 6.92 | 26 | -38 | -4 |
|  | L Parietal Operculum | 10.94 | -44 | -28 | 16 |
|  | R Parietal Operculum | 13.08 | 62 | -32 | 26 |
|  | R Brain Stem | 8.75 | 8 | -30 | -6 |
|  | L Brain Stem | 7.47 | -6 | -34 | -44 |
|  | L Middle Temporal Gyrus, posterior | 7.51 | -62 | -32 | 0 |
|  | R Planum Temporale | 8.17 | 40 | -34 | 16 |
|  | R Supramarginal Gyrus, posterior | 18.17 | 54 | -40 | 10 |
|  | L Supramarginal Gyrus, posterior | 11.6 | -56 | -42 | 24 |
|  | L Superior Parietal Lobule | 19.13 | -34 | -44 | 62 |
|  | R Superior Parietal Lobule | 14.83 | 24 | -48 | 68 |
|  | L Temporal Occipital Fusiform Cortex | 18.72 | -42 | -48 | -20 |
|  | R Angular Gyrus | 8.34 | 62 | -50 | 42 |
|  | L Angular Gyrus | 8.25 | -54 | -52 | 46 |
|  | R Temporal Occipital Fusiform Cortex | 16.89 | 44 | -54 | -20 |
|  | R Precuneus | 15.7 | 6 | -54 | 24 |
|  | L Precuneus | 11.39 | -6 | -58 | 42 |
|  | R Lateral Occipital Cortex, superior | 5.97 | 24 | -60 | 64 |
|  | L Lateral Occipital Cortex, superior | 8.87 | -46 | -62 | 48 |
|  | L Lateral Occipital Cortex, inferior | 19.58 | -48 | -66 | 10 |
|  | R Lateral Occipital Cortex, inferior | 20.95 | 50 | -62 | 6 |
|  | R Lingual Gyrus | 15.93 | 12 | -62 | -2 |
|  | L Lingual Gyrus | 22.34 | -6 | -74 | 4 |
|  | R Intracalcarine Cortex | 21.86 | 6 | -70 | 8 |
|  | R Supracalcarine Cortex | 20.60 | 0 | -74 | 12 |
|  | L Supracalcarine Cortex | 21.08 | -2 | -84 | 12 |
|  | L Occipital Fusiform Gyrus | 5.97 | -40 | -74 | -16 |
|  | R Cuneus | 14.09 | 2 | -80 | 30 |

^a^ Region used in EMA analyses. ^c^ Harvard-Oxford right amygdala, *p*=99%. ^c^ Harvard-Oxford left amygdala, *p*=98%.

**Table S4. Relations between frontocortical reactivity to threat anticipation and real-world negative affect, controlling for subcortical reactivity.**

|  | *t* | *β* |
| --- | --- | --- |
| Threat-Anticipation Frontocortical Composite^a^ | -0.37 | -0.02 |
| Stressor (vs. Absent) | 15.12*** | 0.34 |
| Threat-Anticipation Frontocortical Composite^a^ × Stressor | -2.43* | -0.09 |
| Threat-Anticipation Subcortical Composite | 0.60 | 0.02 |
| Threat-Anticipation Subcortical Composite × Stressor | 0.89 | 0.03 |

^a^ The *Composite* term tests relations between frontocortical function and tonic (stressor-independent) negative affect. The *Composite × Stressor* term tests relations between frontocortical function and reactive (stressor-dependent) negative affect. * *p* ≤ 0.05, ** *p* < 0.01, *** *p* < 0.001

***Continued…***

**Table S5. Relations between frontocortical reactivity to threat anticipation and real-world anxiety and depression*.***

|  | Anxiety | |  | Depression | |
| --- | --- | --- | --- | --- | --- |
|  | *t* | *β* |  | *t* | *β* |
| Threat-Anticipation Frontocortical Composite^a^ | -0.26 | -0.01 |  | 0.40 | 0.01 |
| Stressor (vs. Absent) | 12.73*** | 0.34 |  | 12.74*** | 0.35 |
| Threat-Anticipation Frontocortical Composite^a^ × Stressor | -1.95* | -0.07 |  | -2.04* | -0.07 |

^a^ The *Composite* term tests relations between frontocortical function and tonic (stressor-independent) anxiety or depression. The *Composite × Stressor* term tests relations between frontocortical function and reactive (stressor-dependent) anxiety or depression. * *p* ≤ 0.05, ** *p* < 0.01, *** *p* < 0.001

**Table S6. Relations between frontocortical reactivity to threat anticipation and real-world negative affect, controlling for stressor frequency.**

|  | *t* | *β* |
| --- | --- | --- |
| Threat-Anticipation Frontocortical Composite^a^ | 0.02 | 0.00 |
| Stressor (vs. Absent) | 14.83*** | 0.34 |
| Threat-Anticipation Frontocortical Composite^a^ × Stressor | -2.42* | -0.07 |
| Stressor Frequency | 2.62** | 0.06 |

^a^ The *Composite* term tests relations between frontocortical function and tonic (stressor-independent) negative affect. The *Composite × Stressor* term tests relations between frontocortical function and reactive (stressor-dependent) negative affect. * *p* ≤ 0.05, ** *p* < 0.01, *** *p* < 0.001

**Table S7. Relations between frontocortical reactivity to threat anticipation and real-world negative affect, controlling for amygdala reactivity to threat-related faces.**

|  | *t* | *β* |
| --- | --- | --- |
| Threat-Anticipation Frontocortical Composite^a^ | -0.34 | -0.01 |
| Stressor (vs. Absent) | 15.00*** | 0.35 |
| Threat-Anticipation Frontocortical Composite^a^ × Stressor | -2.21* | -0.07 |
| Threat-Related Faces Amygdala Composite | -0.45 | -0.01 |
| Threat-Related Faces Amygdala Composite × Stressor | -1.56 | -0.04 |

^a^ The *Composite* term tests relations between frontocortical function and tonic (stressor-independent) negative affect. The *Composite × Stressor* term tests relations between frontocortical function and reactive (stressor-dependent) negative affect. * *p* ≤ 0.05, ** *p* < 0.01, *** *p* < 0.001

***Continued…***

**Table S8. Relations between frontocortical reactivity to threat anticipation and real-world positive affect*.***

|  | *t* | *β* |
| --- | --- | --- |
| Threat-Anticipation Frontocortical Composite^a^ | 0.95 | 0.05 |
| Positive Event (vs. Absent) | 23.77*** | 0.51 |
| Threat-Anticipation Frontocortical Composite^a^ × Positive Event | -1.46 | -0.04 |

^a^ The *Composite* term tests relations between frontocortical function and tonic (stressor-independent) *positive* affect. The *Composite × Stressor* term tests relations between frontocortical function and reactive (stressor-dependent) *positive* affect. * *p* ≤ 0.05, ** *p* < 0.01, *** *p* < 0.001

**Table S9. Relations between frontocortical reactivity to threat anticipation and real-world negative affect, controlling for individual differences in trait negative emotionality.**

|  | *t* | *β* |
| --- | --- | --- |
| Threat-Anticipation Frontocortical Composite^a^ | 0.18 | 0.01 |
| Stressor (vs. Absent) | 15.87*** | 0.35 |
| Threat-Anticipation Frontocortical Composite^a^ × Stressor | -2.17* | -0.06 |
| Trait Negative Emotionality | 4.21** | 0.10 |
| Trait Negative Emotionality × Stressor | 4.53*** | 0.11 |

^a^ The *Composite* term tests relations between frontocortical function and tonic (stressor-independent) negative affect. The *Composite × Stressor* term tests relations between frontocortical function and reactive (stressor-dependent) negative affect. * *p* ≤ 0.05, ** *p* < 0.01, *** *p* < 0.001

**Table S10. Relations between regional frontocortical reactivity to threat anticipation and real-world negative affect*.***

|  | Threat-Anticipation  dlPFC/FP | |  | Threat-Anticipation  FrO | |  | Threat-Anticipation  AI | |  | Threat-Anticipation  MCC | |
| --- | --- | --- | --- | --- | --- | --- | --- | --- | --- | --- | --- |
|  | *t* | *β* |  | *t* | *β* |  | *t* | *β* |  | *t* | *β* |
| Brain^a^ | 0.53 | 0.02 |  | -0.78 | -0.02 |  | 0.84 | 0.02 |  | -0.59 | -0.02 |
| Stressor (vs. Absent) | 15.02*** | 0.34 |  | 15.14*** | 0.34 |  | 14.98*** | 0.34 |  | 15.17*** | 0.34 |
| Brain^a^ × Stressor | -1.39 | -0.03 |  | -2.56* | -0.06 |  | -1.59 | -0.04 |  | -2.63** | -0.07 |

^a^ Focal neural metric. The *Brain* term tests relations between regional brain function and tonic (stressor-independent) negative affect. The *Composite × Stressor* term tests relations between regional brain function and reactive (stressor-dependent) negative affect. * *p* ≤ 0.05, ** *p* < 0.01, *** *p* < 0.001

***Continued…***

**Table S11. Relations between MCC reactivity to threat anticipation and real-world negative affect, controlling for AI and dlPFC/FP reactivity.**

|  | *t* | *β* |
| --- | --- | --- |
| MCC Threat-Anticipation^a^ | -1.76 | -0.07 |
| Stressor (vs. Absent) | 15.06*** | 0.35 |
| MCC Threat-Anticipation^a^ × Stressor | -2.05* | -0.08 |
| AI Threat-Anticipation | 1.34 | 0.05 |
| AI Threat-Anticipation × Stressor | -0.10 | 0.04 |
| dlPFC Threat-Anticipation | 1.03 | 0.04 |
| dlPFC Threat-Anticipation × Stressor | 0.58 | 0.02 |

^a^ The *MCC* term tests relations between MCC function and tonic (stressor-independent) negative affect. The *MCC × Stressor* term tests relations between MCC function and reactive (stressor-dependent) negative affect. * *p* ≤ 0.05, ** *p* < 0.01, *** *p* < 0.001

**Table S12. Relations between FrO reactivity to threat anticipation and real-world negative affect, controlling for AI and dlPFC/FP reactivity.**

|  | *t* | *β* |
| --- | --- | --- |
| FrO Threat-Anticipation^a^ | -1.96* | -0.08 |
| Stressor (vs. Absent) | 15.05*** | 0.34 |
| FrO Threat-Anticipation^a^ × Stressor | -1.94* | -0.07 |
| AI Threat-Anticipation | 1.54 | 0.06 |
| AI Threat-Anticipation × Stressor | 0.13 | 0.00 |
| dlPFC Threat-Anticipation | 0.39 | 0.03 |
| dlPFC Threat-Anticipation × Stressor | 0.80 | 0.01 |

^a^ The *FrO* term tests relations between FrO function and tonic (stressor-independent) negative affect. The *FrO × Stressor* term tests relations between FrO function and reactive (stressor-dependent) negative affect. * *p* ≤ 0.05, ** *p* < 0.01, *** *p* < 0.001

**Table S13. Relations between MCC and FrO reactivity to threat anticipation and real-world negative affect, controlling for the other region.**

|  | *t* | *β* |
| --- | --- | --- |
| MCC Threat-Anticipation^a^ | -0.04 | -0.00 |
| Stressor (vs. Absent) | 15.14 | 0.34 |
| MCC Threat-Anticipation^a^ × Stressor | -1.07 | -0.04 |
| FrO Threat-Anticipation^b^ | -0.51 | -0.02 |
| FrO Threat-Anticipation^b^ × Stressor | -0.95 | -0.03 |

^a^ The *MCC* term tests relations between MCC function and tonic (stressor-independent) negative affect. The *MCC × Stressor* term tests relations between MCC function and reactive (stressor-dependent) negative affect. ^b^ The *FrO* term tests relations between FrO function and tonic (stressor-independent) negative affect. The *FrO × Stressor* term tests relations between FrO function and reactive (stressor-dependent) negative affect. * *p* ≤ 0.05, ** *p* < 0.01, *** *p* < 0.001

***Continued…***

**Supplementary Method for Meta-Analytic Analyses of the Cingulo-Opercular Circuit**

Neurosynth—a cloud-based suite of neuroinformatics tools and databases—was used to clarify the functional architecture of regions highlighted by our Brain-EMA results (Yarkoni, Poldrack, Nichols, Van Essen, & Wager, 2011).

**Functional connectivity.** Intrinsic functional connectivity was assessed using an automated seed-based approach and data from the Yeo-Buckner database, which incorporates ‘resting-state’ fMRI data from 1,000 participants (Yeo et al., 2011). For illustrative purposes, connectivity maps were arbitrarily thresholded at a conservative level (*p* < 1.76 × 10^-10^, uncorrected).

**Meta-analytic co-activation.** Regional co-activation patterns were meta-analytically assessed using an automated seed-based approach and a database of >500,000 stereotactic coordinates from >14,000 published imaging studies. Meta-analytic co-activation maps were thresholded using FDR *q* < .01 (whole-brain corrected).

**Meta-Analytic Results**

The present results raise the possibility that the MCC and FrO represent a meaningful functional circuit. If so, then we would expect them to show robust intrinsic functional connectivity in the absence of an explicit task *and* a consistent pattern of co-activation across experimental challenges (Laird et al., 2013; Yeo et al., 2011). We used Neurosynth—a cloud-based suite of neuroinformatics tools and databases—to test these two predictions (Yarkoni et al., 2011). We began by assessing the intrinsic functional connectivity of the MCC and FrO in the Yeo-Buckner database (n = 1,000; Yeo et al., 2011), using the peak locations identified in the threat-anticipation task as seeds (cf. **Figure 2c** in the main report). In each case, there was substantial functional connectivity with the other seed locations (e.g., right MCC ↔ left FrO; *p* < 1.76 × 10^-10^, uncorrected; **Supplementary** **Figure S3a**).

We used a conceptually similar seed-based approach to probe MCC-FrO co-activation. This analysis leveraged a computer-generated database of >500,000 stereotactic coordinates derived from >14,000 published imaging studies. This allowed us to perform a series of automated meta-analyses, each quantifying the likelihood that activation in any one of the seed locations is associated with significant co-activation in the other three (FDR *q* < .01, whole-brain corrected). Mirroring the functional connectivity results, this revealed robust co-activation across the four seed locations (Supplementary **Figure S3b**).

Taken together, these results provide clear evidence that the MCC and FrO form a cingulo-opercular circuit that is sensitive to threat-anticipation in the laboratory and associated with dampened stressor reactivity in the real world.

***Continued…***

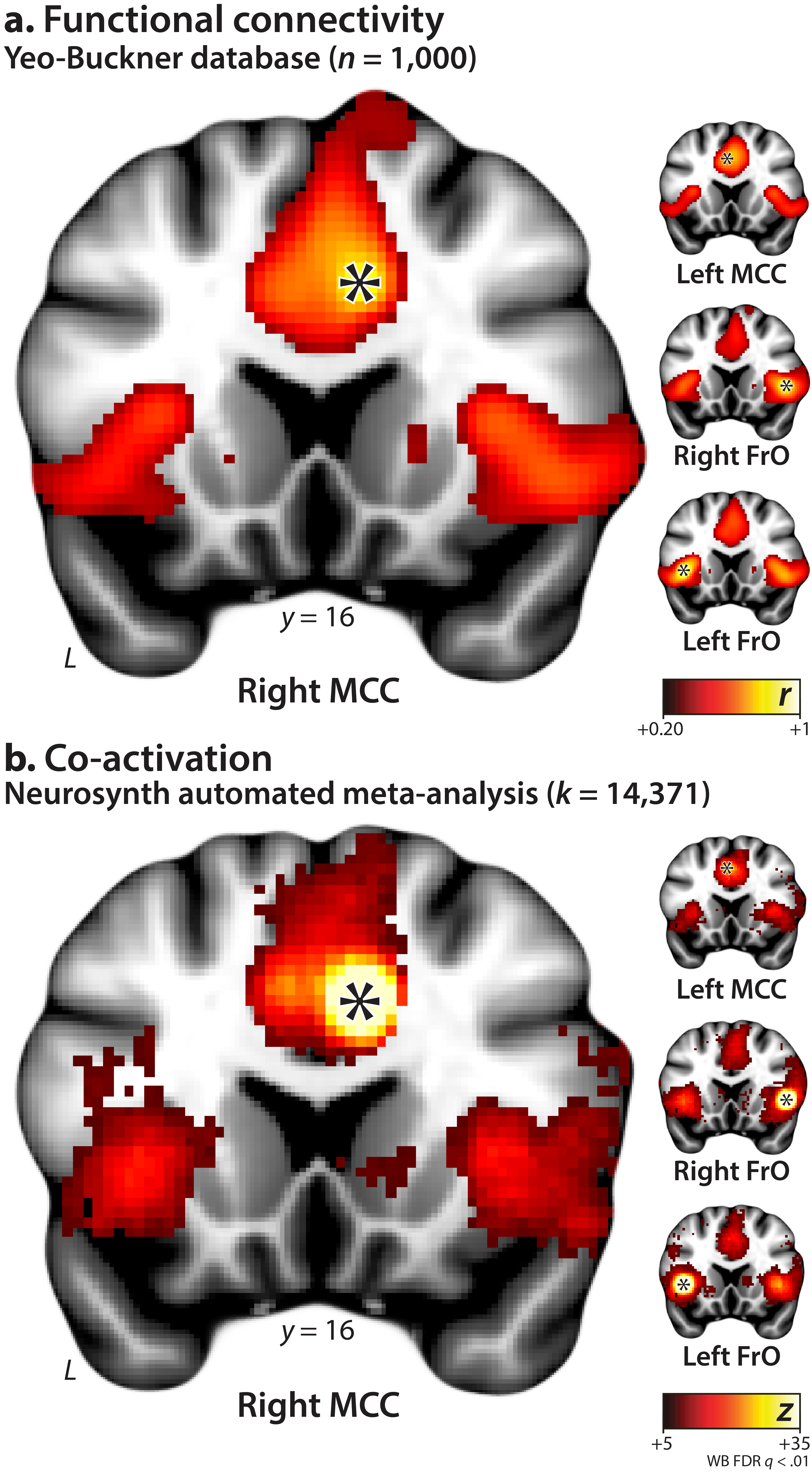
**Supplementary Figure S3**. **The MCC and FrO form a coherent functional neuroanatomical circuit*.*** Asterisks depict the approximate location of each seed. **a. Functional connectivity.** Panel depicts the results of seed-based analyses of intrinsic functional connectivity between the MCC and FrO. Analyses were performed using Neurosynth and the Yeo-Buckner database of 1,000 ‘resting’ fMRI assessments. See **Supplementary Table S1** for seed coordinates. Connectivity maps are shown using an arbitrary threshold (*p* < 1.76 × 10^-10^, uncorrected). **b.** **Co-activation.** Panel depicts the results of seed-based co-activation meta-analyses. Meta-analyses were automatically computed using the Neurosynth database, which encompasses peak coordinates from >15,000 published neuroimaging studies. Co-activation maps are depicted using a FDR *q* < .01, whole-brain corrected threshold. **Abbreviations**—FDR, false discovery rate; FrO, frontal operculum; *k*, the number of studies used in the meta-analysis; L, left; MCC, midcingulate cortex; WB, whole-brain corrected.

***Continued…***
